# Dietary Non-protein Amino Acid AZE as an Environmental Trigger Activating Latent Genetic Susceptibility in *C9orf72*-ALS Pathogenesis

**DOI:** 10.64898/2026.01.12.697838

**Authors:** Chongjiu Chen, Zhanyun Lv, Guangyu Cui, Hui Gong, Jianfeng Hua, Tao Luo, Hongyun Bi, Xinrui Zhang, Qinqin Cui, Hongying Zhu, Xing Guo, Nengyin Sheng, Chengyong Shen, Dongsheng Fan, Ke Zhang, Huaqing Liu, Ge Bai

## Abstract

In human disease, clinical manifestations are not always tightly correlated with genetic etiology. While the GGGGCC (G4C2) hexanucleotide repeat expansion in the *C9orf72* gene is the most prevalent genetic cause of ALS, a subset of mutation carriers stays asymptomatic. The molecular mechanisms underlying this incomplete penetrance remain elusive. Here, we identify L-azetidine-2-carboxylic acid (AZE), a dietary non-protein amino acid, as a critical environmental risk factor capable of unmasking latent genetic susceptibility in C9orf72-ALS pathogenesis. Specifically, AZE disrupts the proteasome-mediated clearance of toxic poly-GA dipeptide repeats derived from G4C2 repeats, triggering pronounced neuroinflammation and motor defects in a C9orf72-ALS mouse model that otherwise exhibits minimal phenotypes. Notably, AZE is widely present in the food chain (particularly in sugar beet) and consistently detectable in the postmortem human brains. A significant positive correlation was observed between sugar beet consumption and the incidence of sporadic C9-ALS cases in European nations. Critically, dietary intake of sugar beet juice is sufficient to activate subclinical genetic susceptibility in C9orf72-ALS mice. These findings reveal a direct diet-gene interplay in governing C9orf72-ALS penetrance, and propose dietary restriction of AZE-rich foods as a tangible strategy for genetically susceptible individuals to potentially delay or mitigate this currently incurable disorder.

## Introduction

While decades of research have linked numerous diseases to genetic mutations, clinical symptoms are not always tightly associated with genetic etiology. Even among individuals carrying identical pathogenic mutations, disease phenotypes can vary remarkably in severity and progression. However, the critical determinants governing the conversion from genetic susceptibility to clinical manifestation remain poorly understood. Elucidating these factors would provide crucial insights for developing preventive strategies and therapeutic interventions.

This genotype-phenotype discordance is particularly well exemplified in amyotrophic lateral sclerosis (ALS), a fatal neurodegenerative disease primarily affecting motor neurons, clinically characterized by muscle atrophy and progressive loss of motor function^1,2^. To date, over 50 genes have been linked to ALS pathogenesis. Among them, the GGGGCC (G4C2) hexanucleotide repeat expansion in *C9orf72* gene represents the most prevalent genetic cause, accounting for approximately 40% of familial and 5% of sporadic cases in European descent, with its incidence varying across different countries^3–5^. Notably, this mutation exhibits incomplete penetrance, with a subset of mutation carriers exhibiting minimal or subclinical phenotypes. The molecular basis underlying this striking variability remains an unsolved puzzle^6,7^.

Important clues to this phenomenon may derive from mechanistic studies of C9-ALS pathogenesis. Emerging evidence has suggested that G4C2 repeat expansions undergo repeat-associated non-ATG (RAN) translation, producing five potentially cytotoxic dipeptide repeats (DPRs): poly-GA, poly-GP, poly-GR, poly-PA, and poly-PR^6,8–12^. Among these, poly-GA is the most abundant in the brain tissues of ALS patients and exhibits a strong propensity to form protein aggregates^13,14^. Overexpression of poly-GA in mice via viral vectors leads to pronounced cytotoxicity and severe neuroinflammation^15^. However, similar phenotypes are not as evident in *C9orf72*-humanized mouse models (C9-ALS mice), which carry a patient-derived bacterial artificial chromosome (BAC) transgene and exhibit substantially lower DPR expression^16–22^. This phenotypic discrepancy likely stems from the differential DPR expression levels between these two experimental systems. Extending this observation to the clinical setting, variability in DPR expression may similarly contribute to the heterogeneous disease penetrance observed in C9-ALS patients.

In contrast to the rapid progress in identifying genetic factors as mentioned above, research on environmental contributors to ALS has lagged behind despite their potential to modulate disease manifestation^23–25^. In this study, we identified L-azetidine-2-carboxylic acid (AZE)—a widely distributed non-protein amino acid (NPAA) in the food chain—as a critical environmental trigger that converts genetic susceptibility into clinical manifestation, thereby modulating disease penetrance in C9-ALS. Mechanistically, AZE disturbs the ubiquitin-proteasome system (UPS) by inducing proteasome overload, leading to the abnormal accumulation of toxic poly-GA dipeptide repeats. Critically, we reveal that the differential incidence of sporadic C9-ALS cases in European nations significantly correlates with national sugar-beet consumption, the richest dietary source of AZE. Furthermore, dietary intake of sugar beet juice is sufficient to activate subclinical genetic susceptibility in a C9-ALS mouse model. Together, these findings establish a mechanistic framework for understanding how latent genetic susceptibility is unlocked by dietary environmental triggers, providing an explanation for the epidemiological variation of C9-ALS across populations. From a translational perspective, our findings suggest that individuals with genetic susceptibility may benefit from avoiding AZE-rich diets. This dietary modification could offer a tangible approach to lower disease penetrance, representing a new perspective for the prevention and management of neurogenetic disorders.

## Results

### AZE selectively unmasks the latent vulnerability in C9-ALS neurons

NPAAs comprise a broad class of compounds distinct from the 20 canonical amino acids that constitute proteins. Often produced as metabolic byproducts by plants and microbes, NPAAs display diverse physicochemical properties and biological activities^26^. Given their documented presence in the diet and potential association with diseases, they have emerged as a class of potential environmental risk factors warranting further investigation^27^. In this study, we explored whether specific NPAAs may contribute to ALS pathogenesis, with particular focus on the most prevalent genetic subtype, C9-ALS.

Our initial screen in NSC34 motor neuron cell lines revealed that most tested NPAAs exhibited minimal cytotoxicity, with several notable exceptions including L-azetidine-2-carboxylic acid (AZE), p-fluorophenylalanine (pFPhe), m-tyrosine (mTyr), L-3,4-dihydroxyphenylalanine (L-DOPA), and L-canavanine (CAN) (Fig. 1a and Extended Data Fig. 1a,b). β-methylamino-L-alanine (BMAA) was also included into the candidate list due to its historical association with an ALS-like syndrome in Guam, although it demonstrated only negligible toxicity in our assay, consistent with the ongoing controversy regarding its pathogenicity^28^. To investigate the potential influences of these candidate NPAAs on C9-ALS pathogenesis, we assessed their effects on the primary neuron cultures derived from the spinal cords of wild-type (WT) and C9-ALS mice^19^. Notably, while both WT and C9-ALS neurons exhibited comparable viability under normal conditions, AZE exposure selectively exacerbated vulnerability in C9-ALS neurons relative to WT controls. This selective effect was not observed with other tested NPAAs (Fig. 1b and Extended Data Fig. 1c-h). These findings were further validated using motor neurons derived from human iPSCs of healthy individuals and C9-ALS patients, where AZE similarly triggered selective vulnerability in C9-ALS motor neurons (Extended Data Fig. 1i).

**Fig. 1.**
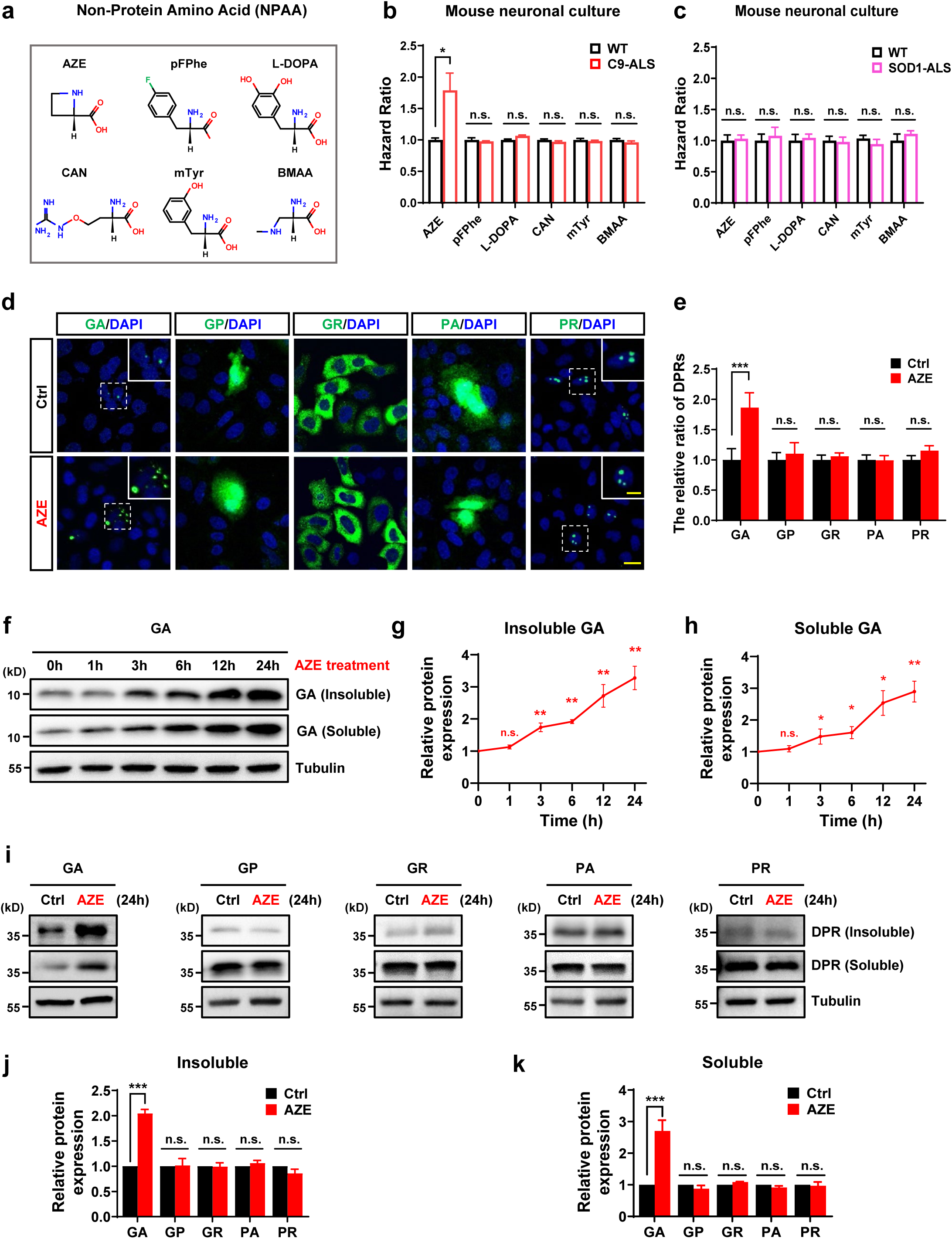
AZE treatment induces the increase of poly-GA levels. a, Chemical structures of non-protein amino acids (NPAAs) identified in a cytotoxicity screening assay. Note that five compounds exhibited greater cytotoxicity compared to the reference compound, BMAA. **b**, Hazard ratio of NPAAs assessed in primary cultured neurons isolated from the spinal cords of wild-type (WT) and *C9orf72*-humanized mouse models (C9-ALS). Note that C9-ALS neurons exhibited increased sensitivity to AZE-induced neurotoxicity compared to WT neurons, while this selective vulnerability was not observed with the other tested NPAAs (AZE, 187μM; pFPhe, 20μM; mTyr, 320μM; L-DOPA, 10μM; CAN, 500μM; BMAA, 1600μM; 24-hour treatment). Note that NPAA concentrations were selected based on dose-toxicity curves where cell viability was maintained at approximately 90%. n ≥ 3 biological replicates. **c**, Hazard ratio of NPAAs in primary cultured neurons from spinal cords of WT and SOD1^G93A^ transgenic mice. Note that no selective vulnerability was observed in SOD1-ALS neurons following treatment with NPAAs, including AZE (AZE, 187μM; pFPhe, 20μM; mTyr, 320μM; L-DOPA, 10μM; CAN, 500μM; BMAA, 1600μM; 24-hour treatment). n ≥ 3 biological replicates. **d**, Representative images of dipeptide repeats (DPRs) in HeLa cells transfected with DPR-expressing plasmids: GA(50)-V5, GP(50)-V5, GR(50)-V5, PA(50)-V5, and PR(50)-V5. V5 staining is depicted in green, and DAPI staining is shown in blue. Note that GA(50)-V5 forms dense protein inclusions; GP(50)-V5, GR(50)-V5, and PA(50)-V5 display a diffuse cellular distribution; PR(50)-V5 is located predominantly to nucleoli. Scale bar, 20μm. Inset: higher magnification of the white boxed area. Scale bar, 10μm. **e**, Quantitative analysis of DPR levels, as shown in (**d**), exhibiting the relative fluorescence signals of DPRs in AZE-treated cells compared to control cells. n ≥ 8 non-overlapping fields (600×600 µm^2^). **f-h**, Western blot analysis showing the time course of AZE treatment (1000μM) on poly-GA levels in HeLa cells transfected with GA(50)-V5-expressing plasmids. Quantitative analysis was conducted by normalizing both soluble and insoluble protein fractions to Tubulin as a loading control (**g,h**). n ≥ 3 independent experiments. **i-k**, Western blot analysis showing the effect of AZE treatment (1000μM, 24-hour treatment) on DPR levels in HeLa cells transfected with DPR-expressing plasmids: GA(50)-GFP, GP(50)-GFP, GR(50)-GFP, PA(50)-GFP, PR(50)-GFP. Quantitative analysis was conducted by normalizing both soluble and insoluble protein fractions to Tubulin as a loading control (**j,k**). n ≥ 3 independent experiments. Note that AZE significantly increased the poly-GA levels but had no detectable effect on other DPRs. Data in **b,c,e,g,h,j,k** are represented as mean ± SEM. Student’s *t* test, *p < 0.05; **p < 0.01; n.s., not significant. See also Extended Data Fig. 1 and Fig. 2.

To further determine whether AZE may affect neuronal viability in other ALS-associated mutations, we tested primary neurons from ALS mice carrying the SOD1^G93A^ mutation (SOD1-ALS). Interestingly, at the same concentration, AZE did not induce similar vulnerability in SOD1-ALS neurons relative to WT controls (Fig. 1c), suggesting that, beyond a general neurotoxic effect, AZE exerts an additional selective toxicity in the context of C9orf72 mutations.

### AZE facilitates the accumulation of poly-GA dipeptide repeats

We next sought to understand how AZE exposure triggers the latent vulnerability in C9-ALS neurons. Unlike WT neurons, C9-ALS neurons express various DPRs due to the G4C2 repeat expansion in the *C9orf72* gene. If these toxic DPRs are not efficiently cleared, they may exacerbate the cell vulnerability. To examine the potential impact of AZE on the clearance of distinct DPR species, we constructed a set of five plasmids, each encoding a DPR variant: GA(50), GP(50), GR(50), PA(50), and PR(50). Immunofluorescence quantification revealed that AZE treatment specifically increased poly-GA levels without significantly affecting other DPR species (Fig. 1d,e). Western blot analysis further confirmed that both soluble and insoluble poly-GA levels began to rise after 3 hours of AZE treatment and continued accumulating over time (Fig. 1f-h), whereas other DPR species showed no apparent changes (Fig. 1i-k). Similarly, AZE had no detectable effect on other ALS-associated mutant proteins, such as SOD1^G93A^, FUS^R521C^, and TDP43^M337V^ (Extended Data Fig. 2a-e). This AZE-driven poly-GA accumulation was further validated in motor neuron cultures and spinal cord organoids derived from human iPSCs of C9-ALS patients, where AZE significantly increased endogenous poly-GA levels originating from the G4C2 repeat expansion (Extended Data Fig. 2f-k).

### AZE induces poly-GA accumulation by disrupting the UPS

These observations prompted us to further investigate how AZE specifically induces poly-GA accumulation. Previous studies indicate that poly-GA is primarily degraded via the ubiquitin-proteasome system (UPS), with ubiquitinated HSP70 playing an essential role in targeting poly-GA for subsequent degradation^29^. Consistent with this, even low concentrations of the proteasome inhibitor MG132 significantly increased both soluble and insoluble poly-GA levels (Fig. 2a-c). Conversely, UPS activation with rolipram effectively promoted poly-GA clearance (Fig. 2d-f). Interestingly, similar modulation of UPS activity had little effect on other DPR species or ALS-associated proteins (Extended Data Fig. 3a-d). These data suggest that poly-GA levels are highly sensitive to even mild UPS dysfunction, which likely stems from its uniquely strong propensity to form protein inclusions compared to other DPRs and ALS-linked mutant proteins. These inclusions have been shown to further suppress proteasome activity, creating a vicious cycle that amplifies its sensitivity to UPS impairment (Fig. 1d and Extended Data Fig. 3e)^14,30–32^.

**Fig. 2.**
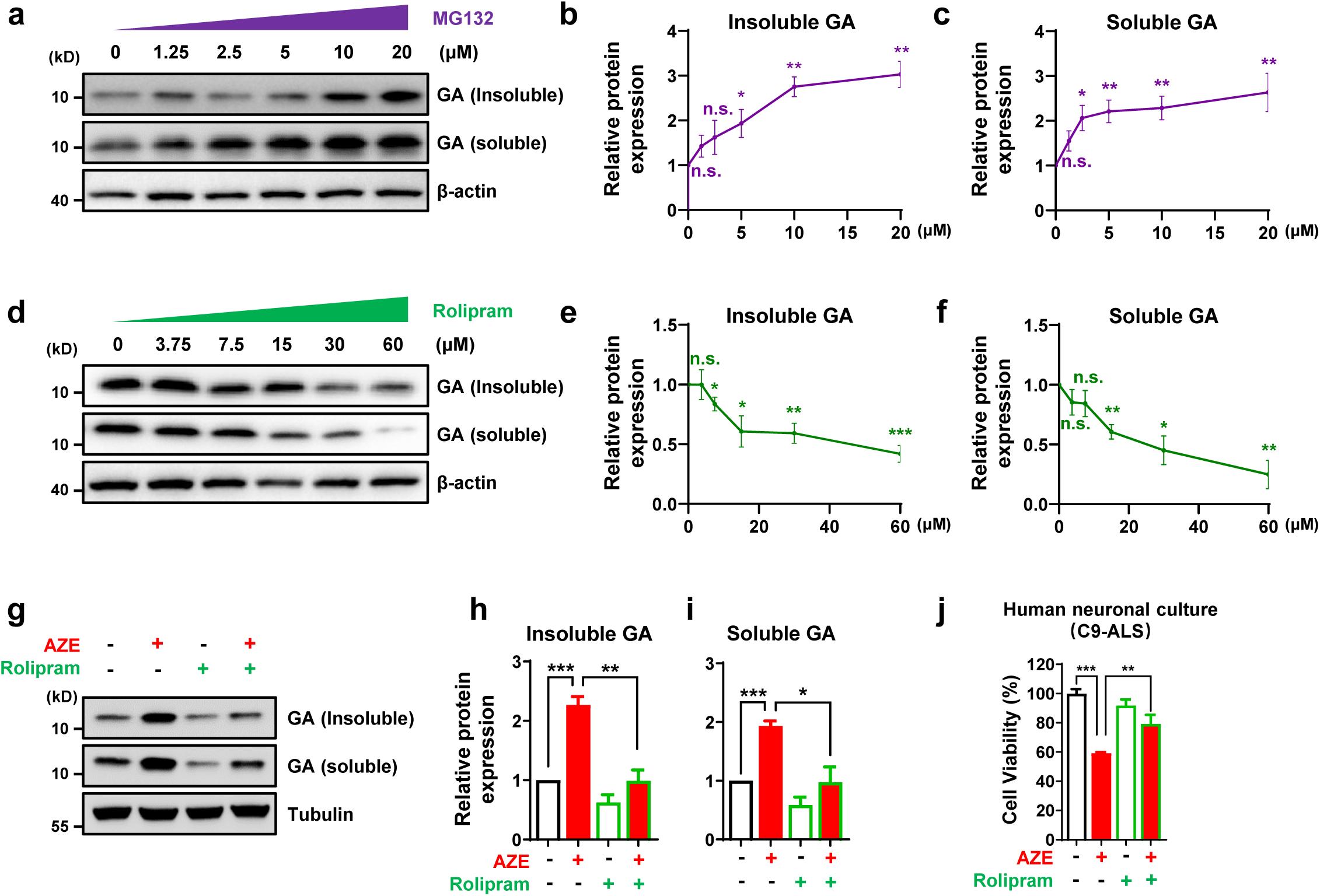
AZE induces poly-GA accumulation by disrupting the UPS. **a-c**, Western blot analysis showing the effect of MG132 treatment (24 hours) on poly-GA levels in HeLa cells transfected with GA(50)-V5-expressing plasmids. Quantitative analysis was conducted by normalizing both soluble and insoluble protein fractions to β-actin as a loading control (**b,c**). n ≥ 3 independent experiments. Note that inhibition of UPS activity by MG132 treatment significantly increased the poly-GA levels. **d-f**, Western blot analysis showing the effect of rolipram treatment (24 hours) on poly-GA levels in HeLa cells transfected with GA(50)-V5-expressing plasmids. Quantitative analysis was conducted by normalizing both soluble and insoluble protein fractions to β-actin as a loading control (**e,f**). n ≥ 3 independent experiments. Rolipram treatment, which enhances UPS activity, significantly decreased the poly-GA levels. **g-i**, Western blot analysis showing that the AZE-induced upregulation of poly-GA levels was diminished by rolipram treatment in HeLa cells transfected with GA(50)-V5-expressing plasmids (AZE, 1000μM; rolipram, 30μM). Quantitative analysis was conducted by normalizing both soluble and insoluble protein fractions to Tubulin as a loading control (**h,i**). n ≥ 3 independent experiments. **j**, Cell viability of iPSC-derived motor neurons from C9-ALS patients following 24-hour treatment with AZE (1000μM) or rolipram (30μM). Note that rolipram treatment attenuated the selective vulnerability of C9-ALS neurons to AZE-induced neurotoxicity. n ≥ 3 biological replicates. Data in **b,c,e,f,h,i,j** are represented as mean ± SEM. Student’s *t* test, *p < 0.05; **p < 0.01; ***p < 0.001; n.s., not significant. See also Extended Data Fig. 3.

To determine whether AZE-induced poly-GA accumulation indeed results from compromised UPS function, we treated AZE-exposed cells with rolipram to enhance the UPS pathway^33,34^. We found that rolipram effectively reversed AZE-induced poly-GA accumulation (Fig. 2g-i and Extended Data Fig. 3f-h). Concurrently, rolipram treatment also attenuated AZE’s toxicity towards C9-ALS neurons (Fig. 2j), suggesting that its toxicity is linked to accumulation of toxic poly-GA in neurons.

### AZE-induced ERAD overwhelms the proteasome to impair poly-GA clearance

Next, we wondered how AZE impairs UPS function. AZE is a known ER stress inducer that activates the ER-associated degradation (ERAD) pathway^35,36^. Given that poly-GA degradation heavily depends on proteasome activity, we propose that AZE-induced activation of ERAD overwhelms the proteasome, thereby reducing poly-GA clearance efficiency and leading to its accumulation (Fig. 3a,b). Consistent with this model, AZE treatment specifically activated the ERAD pathway, accompanied by increased ubiquitinated protein levels, while such effect was not observed with other tested NPAAs (Fig. 3c-e), which also failed to trigger poly-GA accumulation (Fig. 3f-h).

**Fig. 3.**
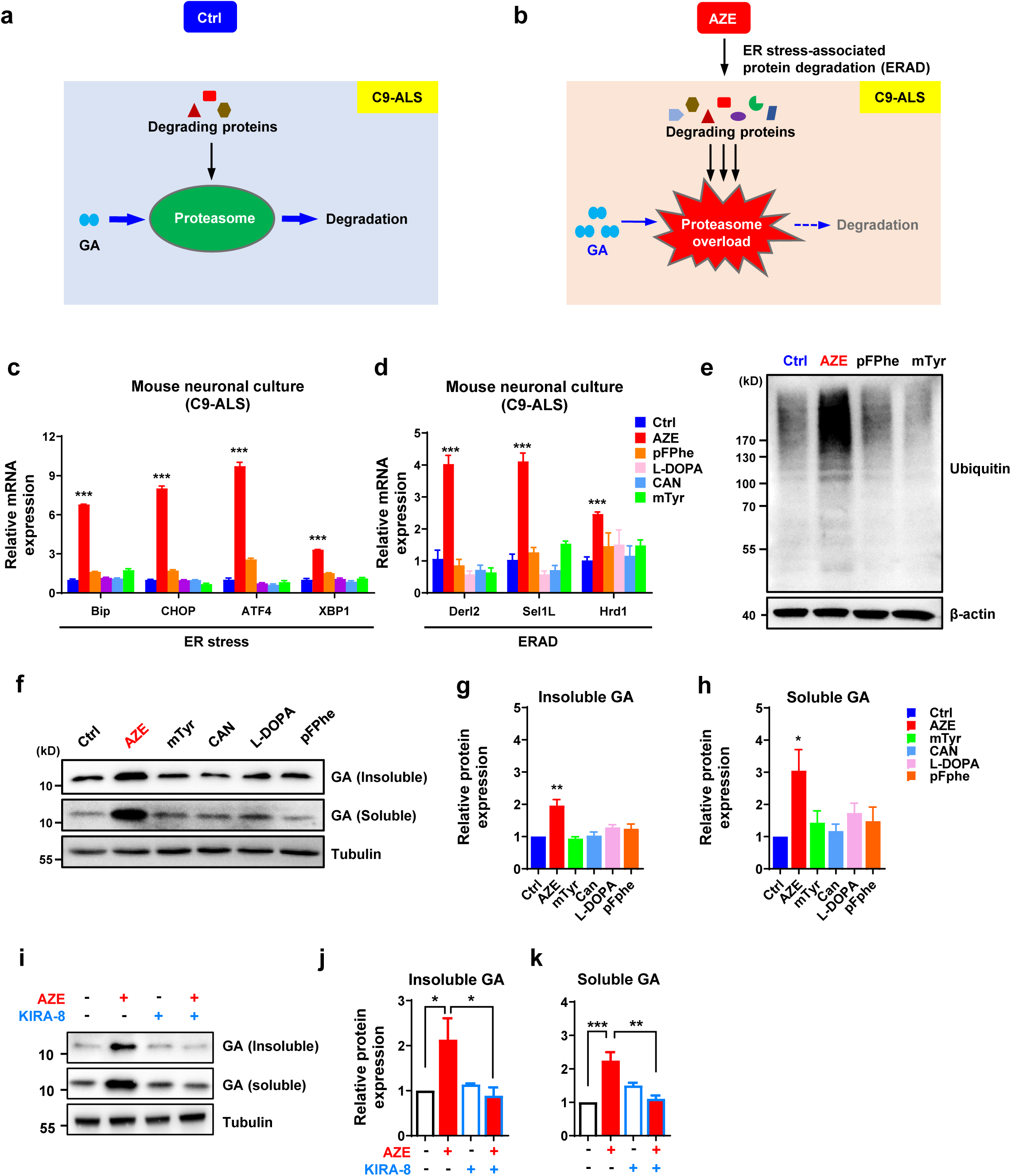
AZE-induced ERAD overwhelms the proteasome. **a**,**b**, Schematic diagram of the proposed mechanism by which AZE increases poly-GA levels in cells. In C9-ALS cells, poly-GA derived from G4C2 repeats relies heavily on the ubiquitin-proteasome system (UPS) for degradation (**a**). However, AZE-induced activation of ER stress-associated protein degradation (ERAD) increases the overall load of proteins targeted for degradation, thereby overwhelming the proteasome. This reduces poly-GA clearance efficiency, leading to its over-accumulation in cells (**b**). **c**,**d**, qPCR analysis showing that AZE treatment exhibits a more robust effect on upregulating ER stress-and ERAD-related genes in motor neuron cultures derived from C9-ALS patient iPSCs compared to other tested NPAAs (AZE, 500μM; pFPhe, 250μM; mTyr, 1000μM; L-DOPA, 100μM; CAN, 50μM; 24-hour treatment). Note that NPAA concentrations were selected based on dose-toxicity curves to ensure comparable effects on cell viability across all tested NPAAs. **e**, Western blot analysis showing increased cellular ubiquitination levels in HeLa cells treated with AZE, but not with other tested NPAAs (AZE, 500μM; pFPhe, 250μM; mTyr, 1000μM; 24-hour treatment). Note that NPAA concentrations were selected based on dose-toxicity curves to ensure comparable effects on cell viability across all tested NPAAs. **f-h**, Western blot analysis showing the effect of NPAAs on poly-GA levels in HeLa cells transfected with GA(50)-V5-expressing plasmids (AZE, 500μM; pFPhe, 250μM; mTyr, 1000μM; L-DOPA, 100μM; CAN, 50μM; 24-hour treatment). Note that NPAA concentrations were selected based on dose-toxicity curves to ensure comparable effects on cell viability across all tested NPAAs. Quantitative analysis was conducted by normalizing both soluble and insoluble protein fractions to Tubulin as a loading control. Note that AZE significantly increased the poly-GA levels, while other tested NPAAs had no significant effect. **i-k**, Western blot analysis showing that the ER stress inhibitor KIRA8 (10nM, 24-hour treatment) attenuates the AZE-induced increase (1000μM, 24-hour treatment) of poly-GA in HeLa cells. Quantitative analysis was conducted by normalizing both soluble and insoluble protein fractions to Tubulin as a loading control. Data in **c,d,g,h,j,k** are represented as mean ± SEM. n ≥ 3 biological replicates. Student’s *t* test, *p < 0.05; **p < 0.01; ***p < 0.001. See also Extended Data Fig. 4.

This mechanism appears both sufficient and necessary for AZE-driven poly-GA accumulation. Indeed, activation of ERAD with thapsigargin (TG) was sufficient to induce poly-GA accumulation (Extended Data Fig. 4a-e). Furthermore, inducing proteasome overload by overexpressing a direct proteasome substrate (GFPodc) also recapitulated the effect of AZE (Extended Data Fig. 4f-h). Conversely, blocking ER stress with KIRA-8 prevented AZE-induced poly-GA accumulation (Fig. 3i-k), confirming the indispensability of this pathway.

### AZE increases poly-GA levels and exacerbates symptoms in C9-ALS mice

Along this line, we sought to further determine whether AZE affects poly-GA levels in C9-ALS mice. To test this possibility, we administered AZE via oral gavage or intraperitoneal injection to WT and C9-ALS mice and subsequently analyzed poly-GA levels in mouse tissues using western blotting. AZE-treated C9-ALS mice showed a significant increase in soluble poly-GA levels across multiple neural tissues, including the spinal cord, cortex, and cerebellum (Fig. 4a-c). The increase was less significant in non-neural tissues such as liver (Fig. 4d). In contrast, poly-GA was nearly undetectable in vehicle-treated C9-ALS mice due to its extremely low expression levels (Fig. 4a-c), consistent with previous reports^19^. Furthermore, this *in vivo* effect of AZE was abolished by ER stress inhibition with KIRA-8, mirroring our *in vitro* findings (Extended Data Fig. 5a-b and Fig. 3i-k). We also did not observe any upregulation of other DPRs upon AZE treatment (data not shown), aligning with its specific effect on poly-GA observed *in vitro* (Fig. 1i-k). Interestingly, neither western blotting nor immunostaining detected insoluble poly-GA in AZE-treated animals (data not shown), possibly because a higher level of poly-GA accumulation is required for its insoluble form to become detectable. By contrast, viral vector-mediated overexpression of poly-GA typically yields both soluble and insoluble forms—an effect that can be further enhanced by AZE exposure (Extended Data Fig. 5c-d).

**Fig. 4.**
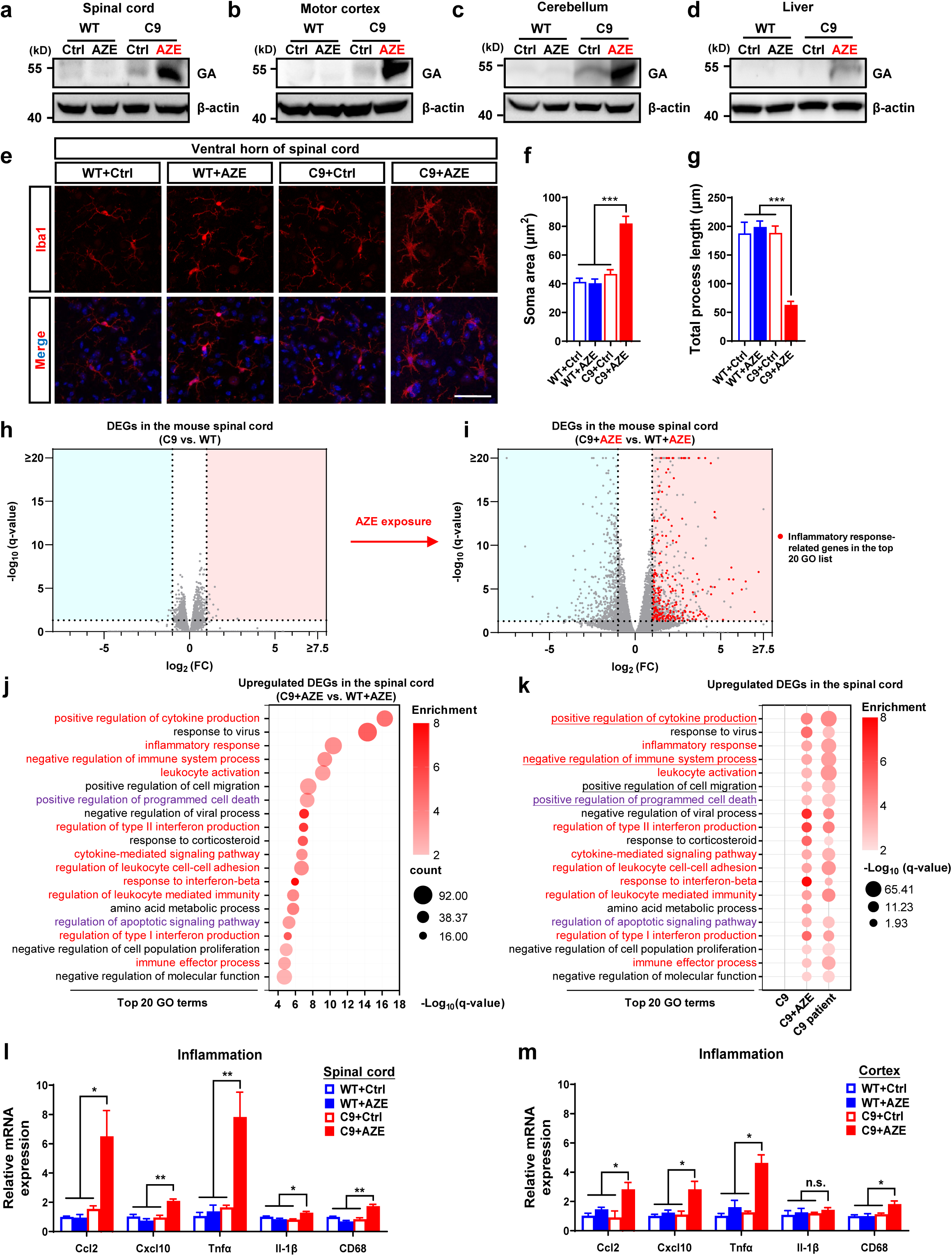
AZE exposure unlocks the latent genetic susceptibility in C9-ALS mice. **a-d**, Western blot analysis showing the effect of AZE treatment (350 mg/kg, i.p., 2 weeks) on poly-GA levels in various neural tissues of wild-type (WT) and *C9orf72*-humanized mouse models (C9-ALS). Control groups were treated with saline (i.p., 2 weeks). Note that a significant increase was observed across multiple neural tissues, while the increase was less significant in non-neural tissues such as the liver. **e**, Representative images of microglia (anti-Iba1, red) in the ventral horn of spinal cords from WT and C9 mice treated with AZE (350 mg/kg, i.p., 4 weeks) or vehicle control (saline, i.p., 4 weeks). Scale bar, 50μm. DAPI staining is shown in blue. Note the significant microglial activation in AZE-treated C9-ALS mice, characterized by enlarged cell bodies and decreased process length. **f,g**, Quantitative analysis of microglia activation in different treatment groups, as shown in (e). n ≥ 3 mice per group. **h**, Volcano plot of RNA-seq data showing minimal differential gene expression in spinal cords of C9-ALS mice vs. wild-type controls. DEGs: differentially expressed genes (fold change ≥ 2; q-value < 0.05). n ≥ 3 mice per group. **i**, Volcano plot of RNA-seq data showing significant differential gene expression in spinal cords of C9-ALS mice vs. wild-type controls following AZE treatment (350 mg/kg, i.p., 4 weeks). Red dots highlight inflammation-related genes in the top 20 enriched Gene Ontology (GO) terms (**j**). n ≥ 3 mice per group. **j**, Bubble plot of Gene Ontology (GO) analysis for genes upregulated in the spinal cords of AZE-treated C9-ALS mice versus AZE-treated WT mice (350 mg/kg, i.p., 4 weeks), as determined by RNA sequencing (fold change ≥ 2; q-value < 0.05). Note that the top 20 enriched GO terms are displayed, with inflammation-related terms highlighted in red and cell death-related terms are highlighted in purple. **k**, Bubble plot comparing enriched GO terms among: C9-ALS mice vs. WT controls, AZE-treated C9-ALS mice vs. AZE-treated WT controls, and human C9-ALS patients vs. healthy controls. Underlined terms represent shared pathways among the top 20 enriched GO terms. Inflammation-related terms are highlighted in red. Cell death-related terms are highlighted in purple. Note that AZE treatment induced upregulation of genes associated with inflammation, cell death and related pathways, recapitulating the characteristic transcriptional signature observed in C9-ALS patients. **l**, qPCR analysis showing that AZE treatment (350 mg/kg, i.p., 4 weeks) significantly induced the upregulation of neuroinflammation-related genes in the spinal cords of C9-ALS mice. n ≥ 3 mice per group. **m**, qPCR analysis showing that AZE treatment (350 mg/kg, i.p., 4 weeks) significantly induced the upregulation of neuroinflammation-related gene expression in the cortex of C9-ALS mice. n ≥ 3 mice per group. Data in **f,g,l,m** are represented as mean ± SEM. One-way ANOVA, *p < 0.05; **p < 0.01; ***p < 0.001; n.s., not significant. See also Extended Data Fig. 5.

Previous studies have indicated that the ectopic expression of poly-GA in the nervous system induces severe inflammatory responses in mice^37^. We wondered whether the AZE-induced poly-GA upregulation was sufficient to trigger a similar response in C9-ALS mice. These mice normally exhibit minimal symptoms, consistent with their low baseline poly-GA levels^19^. Immunofluorescence staining revealed significant microglial activation in multiple brain regions of AZE-treated C9-ALS mice, including the cortex and spinal cord, characterized by enlarged cell bodies and decreased process length (Fig. 4e-g and Extended Data Fig. 5e-g). This phenomenon was absent in both AZE-treated WT mice and vehicle-treated C9-ALS mice (Fig. 4e-g and Extended Data Fig. 5e-g). Consistently, RNA sequencing analysis indicated that AZE treatment triggered the upregulation of numerous inflammatory response-related genes in cortical and spinal cord tissues of C9-ALS mice compared to WT controls, which otherwise exhibit minimal transcriptional differences (Fig. 4h-j, Extended Data Fig. 5h and supplementary Table 1). These results were subsequently validated by quantitative PCR (qPCR) analysis (Fig. 4l,m). Moreover, gene ontology (GO) analysis showed that these molecular changes closely resembled the characteristic transcriptional signatures observed in C9-ALS patients (Fig. 4k and Extended Data Fig. 5i).

Concurrent with these severe inflammatory responses, we observed pronounced motor neuron atrophy in AZE-treated C9-ALS mouse spinal cords (Fig. 5a-c), accompanied by substantial upregulation of cell death-related genes (Fig. 5d). These findings collectively indicate that AZE exposure induced a compromised cellular state in C9-ALS mouse spinal cords, which may contribute to observed neuroinflammation in these mice. However, AZE did not lead to significant neuronal loss or p-TDP43 aggregation in the spinal cord, cortex, or other brain regions of C9-ALS mice (Extended Data Fig. 6a-e and data not shown), phenomena previously reported in viral-mediated poly-GA overexpression models^15^. Interestingly, our findings recapitulate the key pathological features observed in poly-GA transgenic mice with lower poly-GA expression levels compared to viral expression systems^37^, suggesting that the extent of pathological changes may be contingent upon the accumulation levels of poly-GA in neural tissues.

**Fig. 5.**
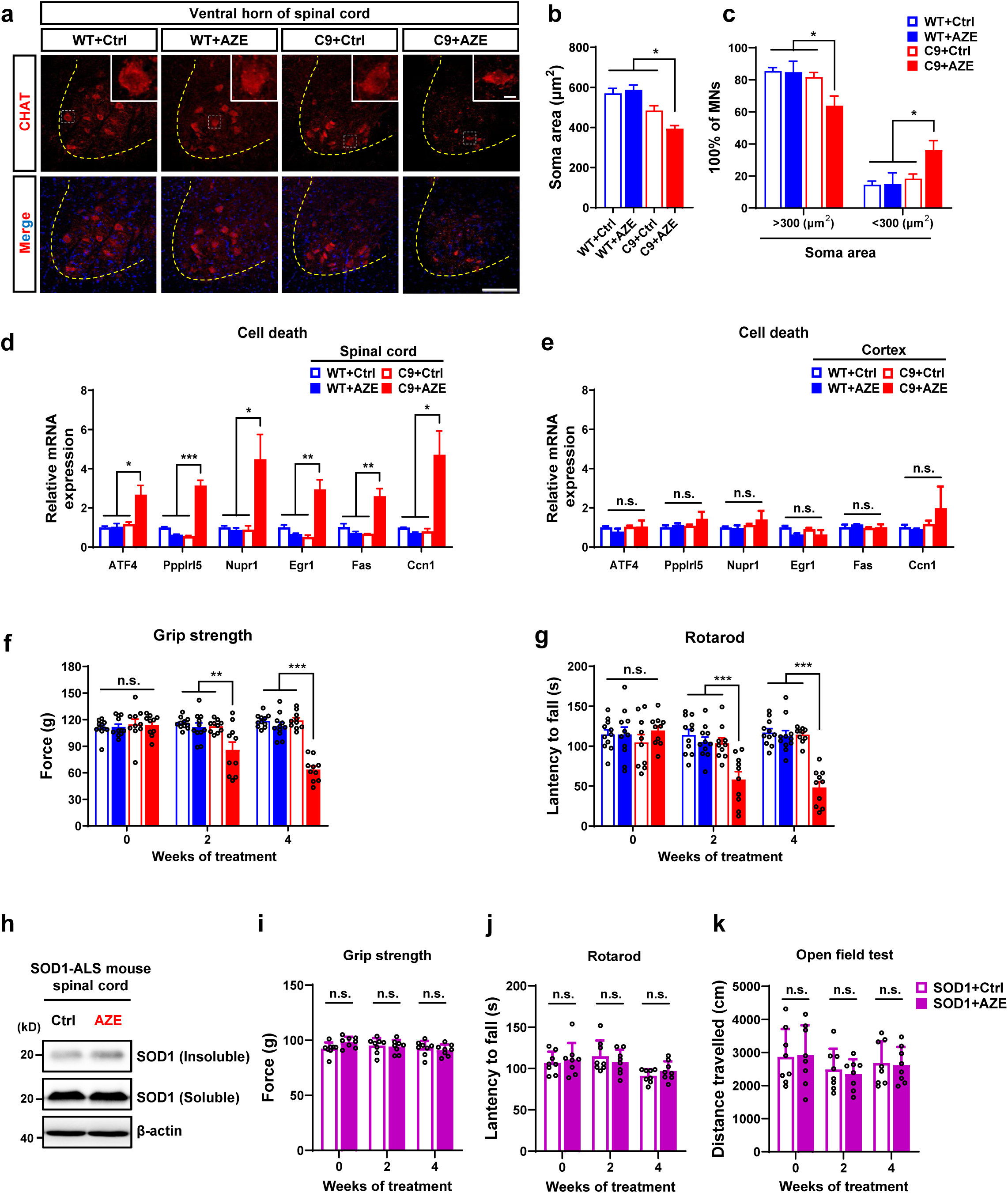
AZE exposure induces motor deficits in C9-ALS mice that otherwise exhibit minimal phenotypes. **a**, Representative images of motor neurons (anti-CHAT, red) in the ventral horn of spinal cords from wild-type (WT) and *C9orf72*-humanized mouse models (C9-ALS) treated with AZE (350 mg/kg, i.p., 4 weeks) or vehicle control (saline, i.p., 4 weeks). DAPI staining is shown in blue. Scale bar, 150μm. Inset: higher magnification of the white boxed area. Scale bar, 10μm. Note the significant reduction of motor neuron size in AZE-treated C9-ALS mice. **b**,**c**, Quantitative analysis of motor neuron size in the ventral horn of spinal cords from different treatment groups, as shown in (**a**). Note that the number of motor neurons with larger size (>300μm^2^) is reduced in AZE-treated C9-ALS mice, while the number of motor neurons with smaller size (<300μm^2^) is correspondingly increased. **d**, qPCR analysis showing that AZE treatment (350 mg/kg, i.p., 4 weeks) significantly induced the upregulation of cell death-related genes in the spinal cords of C9-ALS mice. n ≥ 3 mice per group. **e**, qPCR analysis showing that AZE treatment (350 mg/kg, i.p., 4 weeks) does not significantly upregulate the expression of cell death-related genes in the motor cortex of C9-ALS mice. n ≥ 3 mice per group. **f**,**g**, Grip strength (**f**), rotarod (**g**) of WT or C9-ALS mice treated with AZE (350 mg/kg, i.p.) or saline for different durations. n ≥ 10 mice per group. **h**, Western blot analysis of SOD1 levels in SOD1^G93A^ transgenic mice (SOD1-ALS) treated with AZE (350 mg/kg, i.p., 4 weeks) or vehicle control (saline, i.p., 4 weeks). n ≥ 3 mice per group. **i-k**, Grip strength (**i**), rotarod (**j**) and open field test (**k**) of 2-month-old WT and SOD1-ALS mice treated with AZE (350 mg/kg, i.p.) or vehicle control (saline, i.p.) for different durations. n = 8 mice per group. Data in **b,c,d,e,f,g,i,j,k** are represented as mean ± SEM. One-way ANOVA, *p < 0.05; **p < 0.01; ***p < 0.001; n.s., not significant. See also Extended Data Fig. 6.

Accompanying these pathological changes, AZE-treated C9-ALS mice exhibited a significant decline in motor function compared to C9-ALS controls (Fig. 5f,g and Extended Data Fig. 6f), while cognitive function remained largely unaffected (Extended Data Fig. 6g,h). This motor-specific impairment is aligned with the more severe neuroinflammatory pathology observed in spinal cords compared to brain tissues (Fig. 4e-g,l,m, 5d,e, and Extended Data Fig. 5e-g). Notably, WT mice treated with the same AZE concentration did not exhibit significant behavioral defects (Fig. 5f,g and Extended Data Fig. 6f).

To determine whether AZE exerts similar effects in other ALS models, we tested the classical SOD1^G93A^-ALS mouse model (SOD1-ALS). AZE treatment did not induce an upregulation of SOD1 protein expression in their spinal cords (Fig. 5h), consistent with our *in vitro* findings (Extended Data Fig. 2a). Accordingly, no significant behavioral differences were observed between AZE-treated SOD1-ALS and WT mice (Fig. 5i-k). Moreover, similar treatment with other toxic NPAA also failed to induce a comparable pathogenic effect, arguing against a nonspecific, general neurotoxic mechanism (Extended Data Fig. 6i-k).

### AZE is present in the food chain

NPAAs are widely present in nature, especially in plants and microorganisms, raising the possibility that AZE may enter the human body through food consumption. To test this possibility, we analyzed AZE levels in various common vegetables, fruits, fish, and fermented foods. Our results showed that AZE is detectable in various food samples at markedly variable concentrations, with sugar beet roots identified as the richest source (>25 mg/kg) (Fig. 6a-c). To determine whether dietary AZE is absorbed systemically, we juiced sugar beet roots and orally administered the juice to mice, followed by tissue analysis for AZE content after 24 hours. AZE was detected in multiple organs, with a notable accumulation in the nervous system (Fig. 6d). To further understand the spatial distribution of AZE in the nervous system, we employed imaging mass spectrometry-based spatially resolved metabolite analysis to examine the spinal cords of sugar beet juice-fed mice^38^. The results revealed that AZE primarily accumulates in the gray matter where neuronal cell bodies are located (Fig. 6e).

**Fig. 6.**
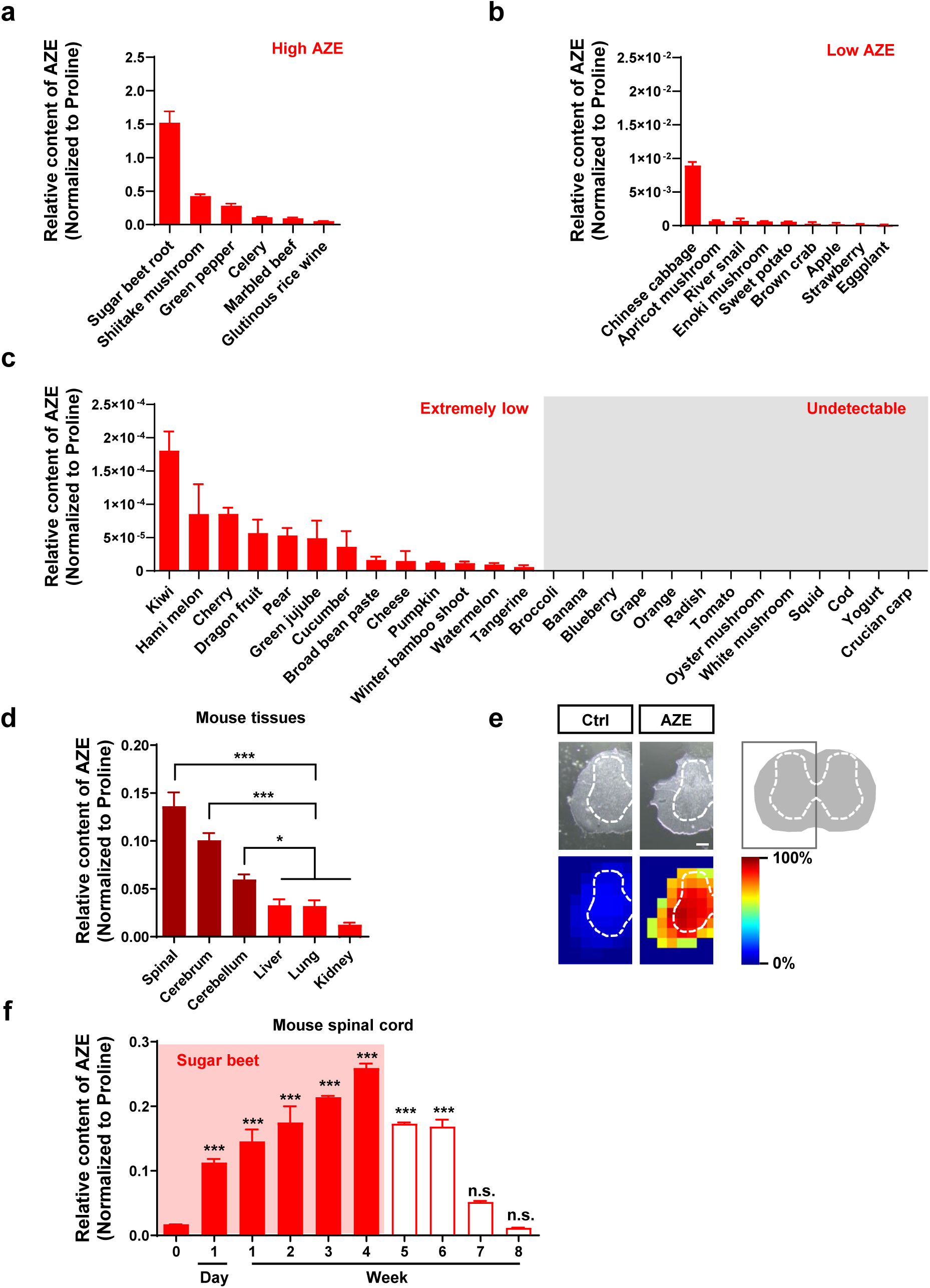
AZE is widely present in the food chain. **a-c**, Bar graph showing the relative abundance of AZE (normalized to proline) in commonly consumed foods. To ensure comparability across species, results are expressed as a ratio of AZE to Proline. **d**, Bar graph showing the relative abundance of AZE (normalized to proline) in various tissues of mice orally administered sugar beet juice (i.g., 24 hours prior to analysis). n ≥ 3 mice per group. **e**, Representative images of AZE distribution in the mouse spinal cord after administered sugar beet juice or controls (i.g., 24 hours prior to analysis) by imaging mass spectrometry. Scale bar, 200μm. **f**, Bar graph showing the relative abundance of AZE (normalized to proline) in the mouse spinal cord at different time points during and after oral administration of sugar beet juice (i.g., daily for up to 4 weeks). n ≥ 3 mice per group. Note that AZE levels increase rapidly upon administration of sugar beet juice but decline gradually following cessation of treatment. Data in **a,b,c,d,f** are represented as mean ± SEM. One-way ANOVA, *p < 0.05; ***p < 0.001; n.s., not significant.

Next, we examined AZE accumulation and metabolic rate *in vivo*. WT mice were subjected to daily intragastric administration of sugar beet juice for one month, and we observed a rapid and continuous accumulation of AZE in the spinal cord (Fig. 6f). Upon cessation of AZE administration, its levels remained stable for approximately two weeks before declining significantly, indicating AZE’s relative stability and potential for accumulation over time (Fig. 6f).

### Sugar beet consumption triggers C9-ALS pathology

Building upon these findings, we further investigated the effects of sugar beet consumption in C9-ALS mice. Our results showed that sugar beet juice administration triggered a significant elevation of poly-GA levels (Fig. 7a-c). This was followed by increased expression of cell death and inflammatory response-related genes, along with microglial activation (Fig. 7d-g), ultimately leading to motor defects (Fig. 7h,i). This entire pathogenic cascade phenocopied the effects of direct AZE administration (∼300mg/kg) (Fig. 4a,e-g, 5d,f, and Extended Data Fig. 6f). Critically, both treatments resulted in comparable spinal cord AZE concentrations (Fig. 7j), demonstrating that the active dietary component, AZE, reaches a pathophysiologically relevant concentration to drive disease in our experimental setting.

**Fig. 7.**
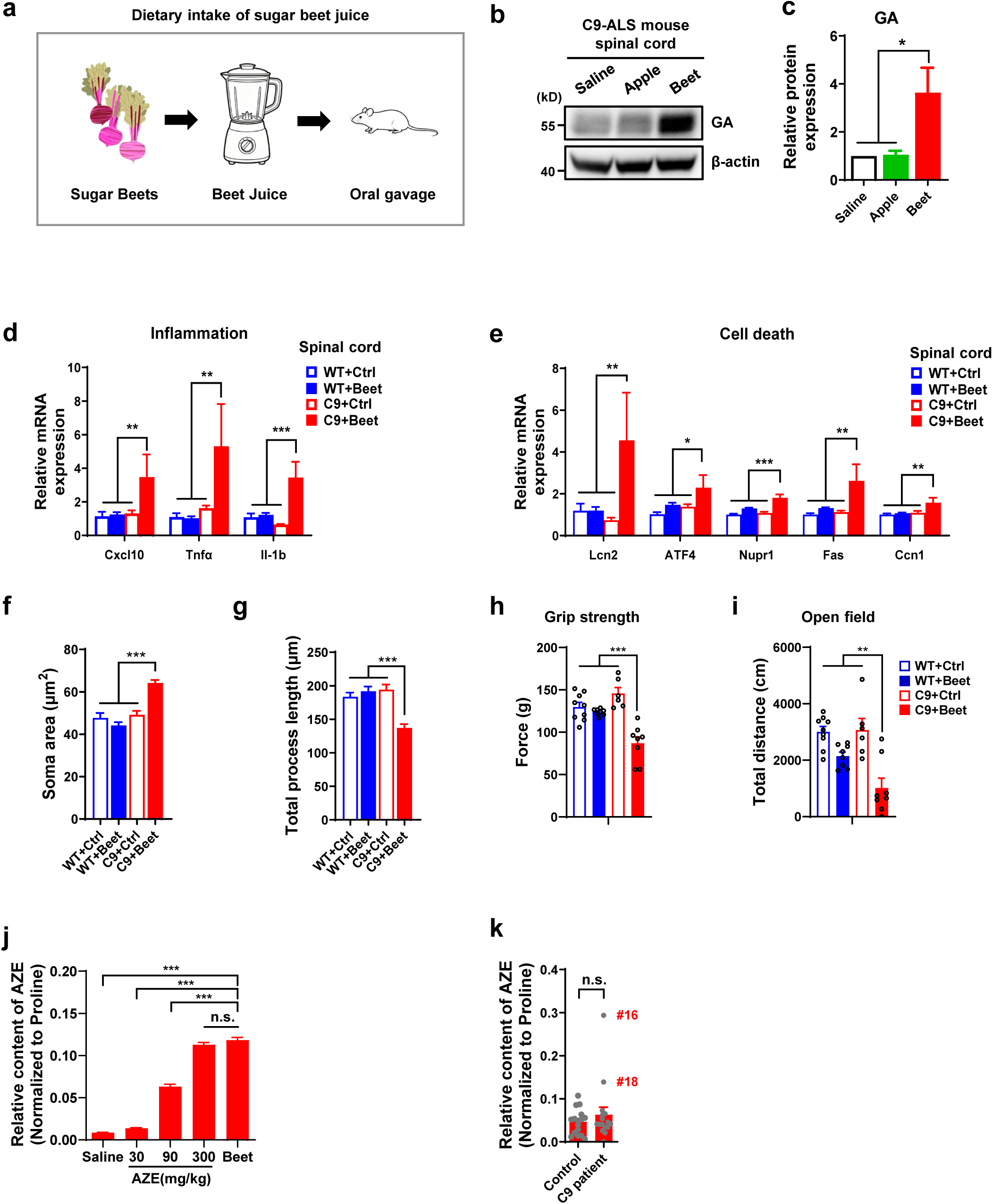
Dietary sugar beet juice triggers neuroinflammation and motor deficits in C9-ALS mice. **a**, Schematic of the sugar beet juice administration in mice. **b**,**c**, Western blot analysis of poly-GA levels in the spinal cord of *C9orf72*-humanized mouse models (C9-ALS) treated with saline, apple juice or sugar beet juice (i.g., 24 hours prior to analysis). Quantitative analysis was conducted by normalizing poly-GA to β-actin as a loading control. Note that sugar beet juice administration induced poly-GA upregulation, while low-AZE-containing foods like apples had no such effect. **d**,**e**, qPCR analysis of the spinal cord from WT or C9-ALS mice after long-term oral gavage with beet juice or saline control (>6 months), showing significant upregulation of neuroinflammation- (**d**) and cell death-related (**e**) genes (n ≥ 3 mice per group). **f**,**g**, Quantification of microglial activation (Iba1+) in the ventral horn of spinal cord from WT or C9-ALS mice after long-term beet juice gavage (>6 months) (n ≥ 3 mice per group) **h**,**i**, Grip strength (**h**) and open field (**i**) tests in WT and C9-ALS mice after long-term sugar beet juice gavage (>6 months) (n ≥ 6 mice per group). **j**, AZE levels in the mouse spinal cord one day after administration via oral gavage of beet juice or intraperitoneal injection of pure AZE at indicated doses (n=3 mice per group). **k**, The relative abundance (normalized to proline) of AZE in post-mortem neural tissues from C9-ALS patients (n=16) and normal controls (n=16) obtained from the Netherlands Brain Bank. Data in **c,d,e,f,g,h,i,j,k** are represented as mean ± SEM. One-way ANOVA, *p < 0.05; **p < 0.01; ***p < 0.001; n.s., not significant. See also Extended Data Fig. 7.

### AZE is detectable in C9-ALS patients

To explore a potential link between AZE and C9-ALS pathogenesis in patients, we analyzed postmortem neural tissues from C9 patients obtained from the Netherlands Brain Bank. We found that AZE was detectable in nearly all tested patient samples, with concentrations varying across individuals (Fig. 7k and Extended Data Fig. 7a). Notably, the average concentration was comparable to levels found in our sugar beet juice-fed and AZE-injected mice (Fig. 7j). Intriguingly, two patients (#16 and #18) exhibited significantly higher AZE levels than the rest of the cohort (Fig. 7k). Consistent with this finding, neuropathological reports from the Netherlands Brain Bank indicated that these same individuals showed pronounced spinal cord pathology (Supplementary Table 4). Patient #16, in particular, was documented with severe neuronal loss in the anterior horn (Supplementary Table 4). In contrast, most other samples presented with relatively mild spinal cord pathology, often accompanied by more prominent frontotemporal dementia (FTD) features (Supplementary Table 4). These clinical observations suggest that higher AZE levels may be associated with more severe spinal cord neuropathology in C9-ALS. This aligns with our experimental findings in AZE-treated C9 mice, which primarily exhibited spinal cord pathology.

Notably, baseline AZE levels did not differ between C9 patients and controls (Fig. 7k and Extended Data Fig. 7a), suggesting that AZE exposure alone may not be intrinsically pathogenic. Instead, the C9 mutation confers specific vulnerability, as evidenced by the fact that equivalent AZE exposure induced motor deficits specifically in C9-ALS models, but not in WT controls (Figure 4,5 and Extended Data Fig. 5). Together, these clinical and experimental data establish a clear gene-environment interaction, wherein a common dietary factor AZE acts as a pathogenic trigger selectively in genetically susceptible individuals.

### Sugar beet consumption correlates with sporadic C9-ALS incidence in Europe

Building on the mechanistic link between AZE and C9-ALS pathology, we sought ecological evidence linking the primary dietary source of AZE—sugar beet—to disease epidemiology. We mainly focus on European nations, which are the world’s primary sugar beet production region with relatively homogeneous demographics, geography, and reliable ALS registry data. A significant positive correlation was observed between per-capita sugar beet consumption and the proportion of sporadic C9-ALS cases, potentially indicative of its important role in increasing disease penetrance (Fig. 8a-e). In contrast, no such correlation was observed between per-capita sugar beet consumption and the proportion of SOD1-ALS among sporadic ALS cases (Fig. 8f). This ecological evidence, alongside our experimental and clinical findings, identifies dietary AZE as a potentially key environmental risk factor in C9-ALS pathogenesis. This finding provides an explanation for the epidemiological variation of C9-ALS across nations, highlighting the translational rationale for dietary modification as a preventative strategy for genetically susceptible individuals.

**Fig. 8.**
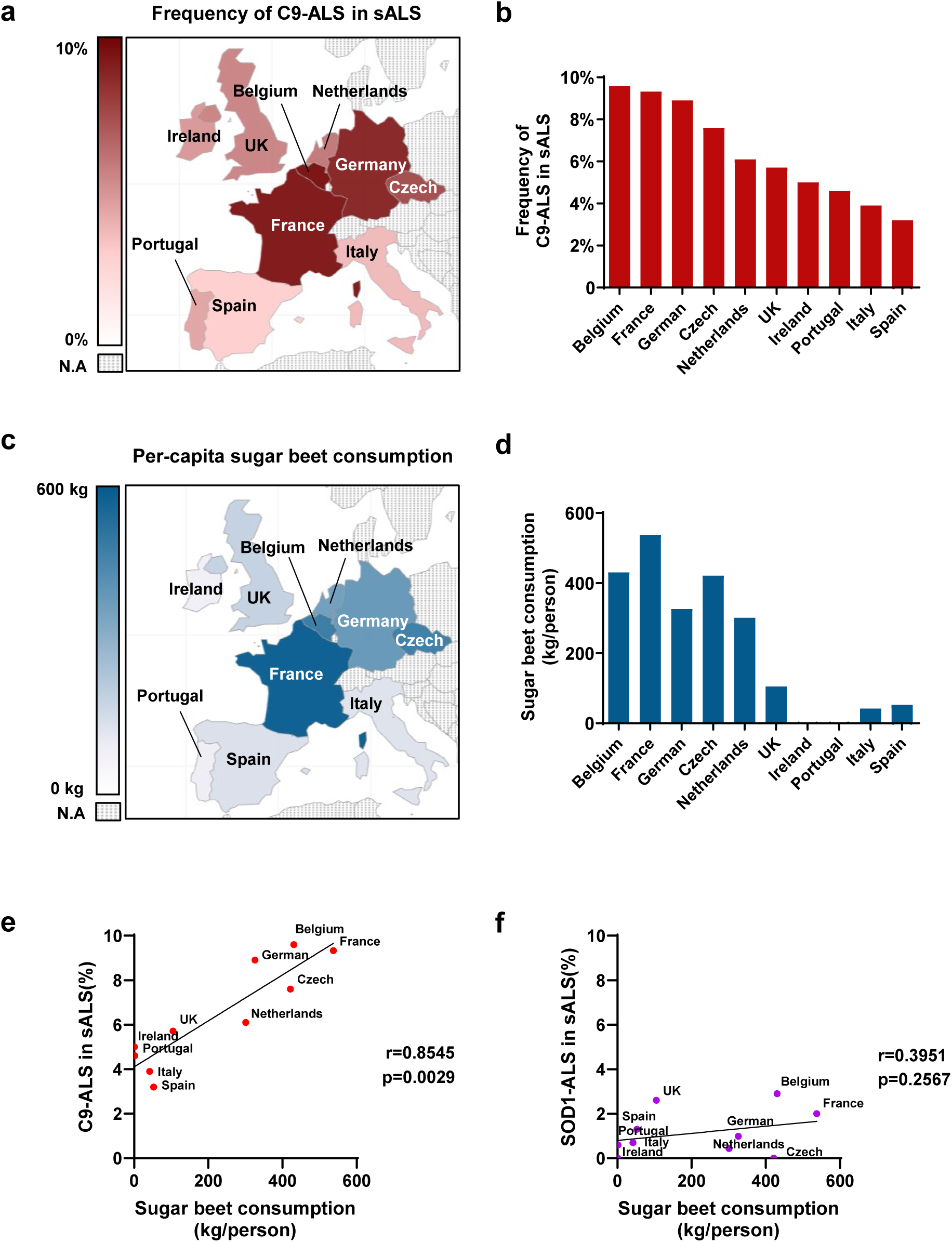
Sugar beet consumption correlates with sporadic C9-ALS incidence in Europe. **a**,**b**, The proportion of C9-ALS cases among sporadic ALS cases in European countries. Countries were included based on geographic proximity and the availability of reliable national ALS registry data. **c**,**d**, Per-capita sugar beet consumption in major European countries, calculated as the total annual amount of sugar beets consumed nationwide for all purposes (direct consumption, processed foods, and livestock feed) divided by population. Given that sugar beet products are widely consumed by humans and are also used in livestock feed, they may enter the human food chain both directly and indirectly (e.g., via. the consumption of marbled beef; see Fig. 6a). Data were obtained from the Food and Agriculture Organization of the United Nations (https://ourworldindata.org/). **e**, Scatter plot showing the significant positive correlation between national per-capita sugar beet consumption and the proportion of C9-ALS among sporadic ALS cases in the analyzed European countries. Spearman’s rank correlation coefficient was used. Note that the sugar beet consumption data correspond to the same calendar year as the sporadic C9-ALS incidence data used for the analysis. See Supplementary Table 4 for details. **f**, Scatter plot showing the absence of a significant correlation between national per-capita sugar beet consumption and the proportion of SOD1-ALS among sporadic ALS cases in the same set of countries. Spearman’s rank correlation coefficient was used. Note that the sugar beet consumption data correspond to the same calendar year as the sporadic SOD1-ALS incidence data used for the analysis. See Supplementary Table 4 for details. See also Extended Data Fig. 7.

## Discussion

### AZE is a key environmental risk factor in C9-ALS pathogenesis

The G4C2 repeat expansion in *C9orf72* is the most prevalent genetic cause of familial ALS^3–5,39^. However, the incomplete penetrance of this mutation and the mild phenotypes observed in *C9orf72*-humanized mouse models strongly suggest that additional factors contribute to disease manifestation^23–25^. Our study identifies AZE, a naturally occurring NPAA present in the food chain, as a potential environmental trigger for C9-ALS pathogenesis. Mechanistically, AZE specifically increases poly-GA levels, inducing neuroinflammation and motor deficits in an otherwise asymptomatic C9-ALS mouse model, demonstrating how environmental exposure can convert genetic risk into clinical manifestation (Fig. 2,3).

As a key step toward establishing AZE as a physiologically relevant trigger, it was crucial to demonstrate that dietary exposure can lead to its accumulation in target tissues at pathologically meaningful levels. Our results showed that AZE is detectable in various foods, with sugar beets identified as its richest source. However, precisely quantifying the absolute concentration of AZE in food is challenging due to its low extraction efficiency. Moreover, the bioavailability of a compound from a natural food matrix (e.g., AZE from beet juice) can differ significantly from that of a purified injection. Therefore, to ensure that our experimental doses were pathophysiologically relevant, we adopted a tissue-level calibration approach (Fig. 6). This calibration confirmed that an i.p. dose of ∼300 mg/kg AZE resulted in spinal cord AZE concentrations similar to those from beet juice gavage. Crucially, this concentration range is also comparable to the levels we detected in the postmortem neural tissues from C9 patients. Since both the ∼300 mg/kg AZE injection and beet juice gavage trigger C9-ALS pathology, these results demonstrates that the dietary intake of AZE can indeed reach a pathophysiologically relevant concentration in neural tissues, thereby faithfully bridging dietary exposure, experimental intervention, and human disease.

Notably, we detected no insoluble poly-GA aggregates or inclusions in AZE-treated C9-ALS mice, suggesting that poly-GA accumulation may need to reach a critical threshold before undergoing the transition into insoluble species (Fig. 4). Nevertheless, the observed neuroinflammation and motor deficits associated with elevated soluble poly-GA levels indicate that inclusion formation is not obligatory to trigger pathological phenotypes. Rather, soluble poly-GA species alone are sufficient to drive C9-ALS pathogenesis, paralleling the neurotoxic effects of soluble Aβ oligomers in Alzheimer’s disease^40^.

In addition, we noticed that AZE-treated C9-ALS mice do not exhibit the full spectrum of ALS phenotypes (Fig. 4,5, Extended Data Fig. 5,6). This partial manifestation may be explained by lack of poly-GA inclusions and relatively low endogenous poly-GA levels compared to exogenous poly-GA overexpression^15,29,37,41^. Moreover, we found that AZE specifically increases poly-GA without affecting other DPR species (Fig. 1). Given that each DPR contributes differently to ALS pathology^15,37,41–48^, this may also account for the limited spectrum of ALS-like symptoms observed in AZE-treated C9-ALS mice.

### Mechanism of AZE-induced poly-GA accumulation

Previous studies have shown that AZE is a proline analog that can be misincorporated into proteins. Given the essential role of proline residues in defining protein backbone structure and overall conformation, its replacement by AZE severely disrupts proper folding^49^. This property confers on AZE a unique ability to potently induce ER stress compared to other NPAAs. Subsequent activation of the ERAD pathway results in an increase of proteins awaiting degradation that overwhelms the proteasome (Fig. 2,3). Notably, compared to other DPRs or ALS-related proteins, poly-GA exhibits unique dependence on the UPS for its degradation and demonstrates exceptional sensitivity to proteasomal dysfunction (Fig. 2). Moreover, even modest accumulation of poly-GA can further inhibit proteasome activity through multiple mechanisms, creating a self-reinforcing pathological cycle that amplifies its sensitivity to UPS impairment^30–32^. This distinctive characteristic explains why compromised proteasome function has the most dramatic effect on poly-GA accumulation^14,30,50^.

### The role of dietary NPAAs in ALS pathogenesis

Various NPAAs are known to be present in food ^51–55^. Epidemiological studies in the 1950s identified an unusually high incidence of a ALS-like symptom among Guam’s Chamorro population, which was initially proposed to be linked to dietary exposure to a non-protein amino acid (NPAA), β-methylamino-L-alanine (BMAA)^52,56,57^. However, the pathogenic role of BMAA lack conclusive experimental support and has remained controversial over the past decades. Moreover, its exposure is geographically restricted. Our study reveals that AZE, but not BMAA and other tested NPAAs, can induce poly-GA accumulation. This may be attributed to the unique ability of AZE to be aberrantly incorporated into proteins, triggering ERAD pathways (Fig. 3). Given the vast diversity of NPAAs, we cannot rule out the possibility that additional ones may contribute to ALS pathogenesis, or synergistic interactions between different NPAAs may occur. Additionally, while AZE appears to have no significant effect on other ALS subtypes such as SOD1^G93A^ (Fig. 5), it remains possible that other untested NPAAs may influence these ALS variants. Exploring these possibilities will be an important direction for future research.

It is important to emphasize that while our study identifies AZE as the first dietary factor clearly influencing C9-ALS pathogenesis, we do not propose AZE as the sole determinant of disease penetrance, given the multifactorial nature of ALS. In fact, our mechanistic findings suggest that other environmental factors affecting ER stress and UPS pathways could potentially contribute to C9-ALS penetrance (Extended Data Fig. 4). The significance of AZE lies in the fact that it is a naturally occurring small molecule compound widely present in foods like sugar beets and can persist long-term in the body. This leads to the possibility of long-term exposure to AZE in large populations. While other ER stress inducers exist in the environment (e.g., heavy metal, viral infections, pesticides), many represent transient or acute stressor affecting limited populations. Additionally, genetic modifiers undoubtedly also play a role in this process. The involvement of these genetic and environmental factors makes C9 penetrance a complex issue that cannot be simply explained by AZE levels alone. For instance, C9-ALS cases occur at a particularly high incidence in Northern Europe compared to other European regions, an observation may reflect founder effects in the population^39,58^. Nevertheless, given that high AZE levels are sufficient to exacerbate C9-ALS pathology and considering the widespread exposure in human populations, the key translational implication of our study is the recommendation of C9-ALS patients to avoid AZE-rich diets. This dietary modification could offer a tangible approach to potentially delay or mitigate this currently incurable disorder. More broadly, it highlights the reduction of disease penetrance by modulating environmental risk factors as a new perspective for the prevention and management of neurogenetic disorders.

## Acknowledgment

We would like to thank Dr. Linhao Ruan for his scientific advice on this project; Dr. Aimin Bao for her assistance with the postmortem human brain sample application process; Dr. Wenyuan Wang for providing iPSCs; Xiaodan Wu for assistance with AZE quantification; members of the Bai lab for discussions on the experiments and manuscript. This work was supported by grants from National Key R&D Program of China (2025YFC3409700 to G.B.), STI2030-Major Projects (2021ZD0202501 to G.B.) and the National Natural Science Foundation of China (82325016, 82150003 and 32421001 to G.B.; 32400780 to H.B.; 32300790 to Q.C.).

## Author contributions

G.B. conceived and supervised the project. C.C. performed biochemical experiments. Z.L., C.C. and X.Z. carried out mouse experiments. G.C. and C.C. carried out immunostaining and imaging. C.C., Z.L., G.C. carried out data analysis. J.H. assisted with primary motor neuron culture. H.G. assisted with iPSC-derived motor neurons culture. T.L. assisted with iPSC-derived spinal cord organoid culture. H.B. assisted with western blot experiments. H.Z. assisted with MALDI-MS imaging. X.G., N.S., C.S., K.Z., D.F. and H.L. contributed to interpretation of results, experimental design, and/or scientific advice. C.C, G.C. and G.B. wrote the primary draft of the manuscript, and all authors contributed to the final version.

## SUPPLEMENTAL INFORMATION

Supplemental information includes 7 figures, and 4 tables.

## Extended Data Figure Legends

**Extended Data Fig. 1.**
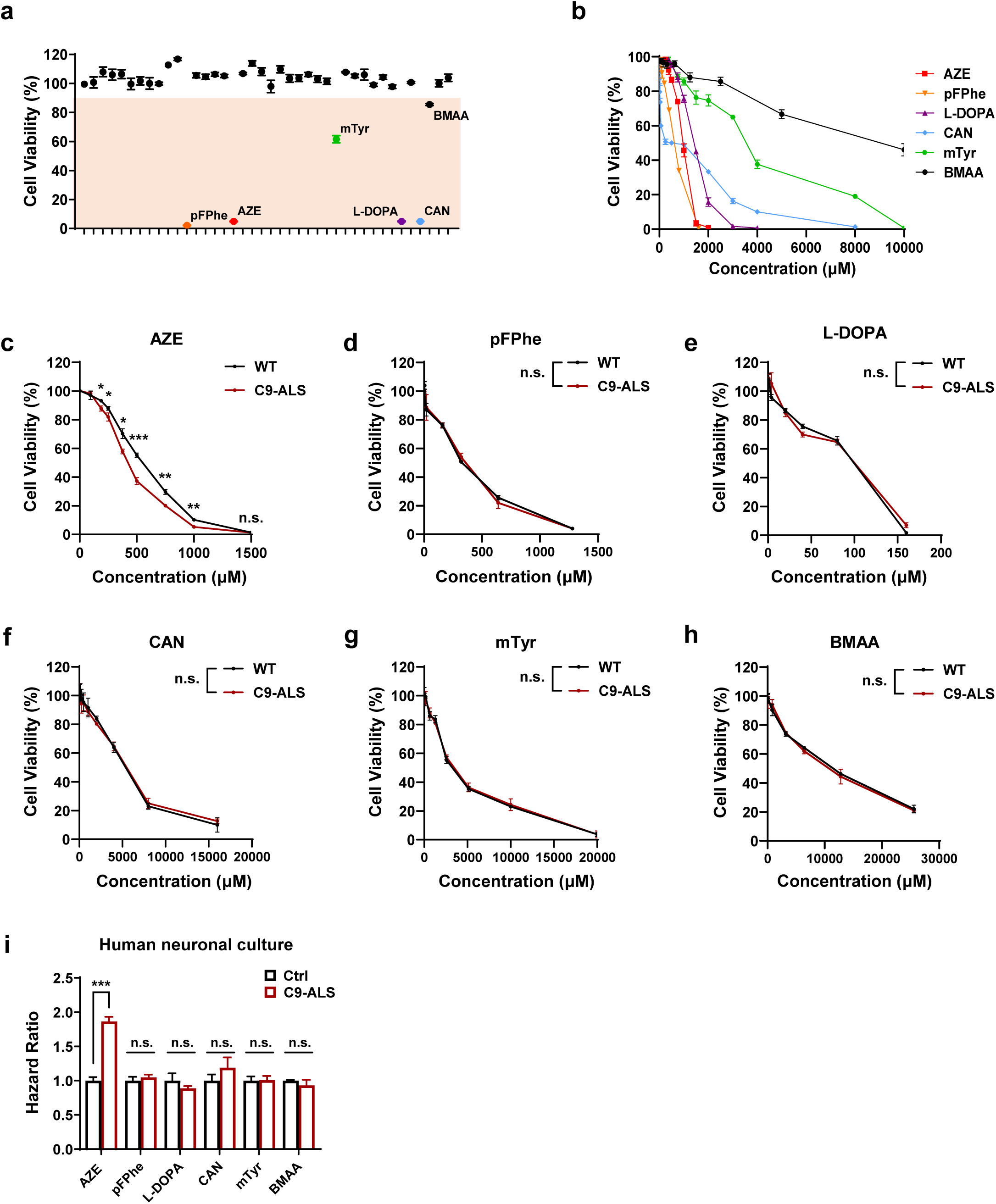
Dose-dependent toxicity of NPAAs in neuronal cultures, related to Fig. 1. **a**, Cell viability assay evaluating the effects of 40 commercially available non-protein amino acids (NPAAs) on NSC34 cells following 96-hour treatment (5mM). Five NPAAs exhibiting toxicity profiles were identified: AZE, pFPhe, mTyr, L-DOPA, CAN. Note that BMAA was included into the candidate list due to its historical association with an ALS-like syndrome in Guam. **b**, Dose-toxicity curves derived from cell viability assays in NSC34 cells treated with varying concentrations of 6 NPAAs for 96 hours. The curves illustrate the relationship between NPAA concentration and cellular toxicity. **c-h**, Dose-toxicity curves of 6 NPAAs assessed in primary cultured neurons isolated from the spinal cords of wild-type (WT) and *C9orf72*-humanized mouse models (C9-ALS). Note that C9-ALS neurons demonstrated increased sensitivity to AZE-induced neurotoxicity compared to WT neurons, while this selective vulnerability was not observed with the other tested NPAAs (24-hour treatment). **i**, Hazard ratio of NPAAs assessed in motor neuron cultures derived from iPSCs of C9-ALS patients and healthy controls. Note that C9-ALS neurons exhibited increased sensitivity to AZE-induced neurotoxicity compared to control neurons, while this selective vulnerability was not observed with the other tested NPAAs (AZE, 250μM; pFPhe, 125μM; mTyr, 250μM; L-DOPA, 12.5μM; CAN, 250μM; BMAA, 1250μM, 24-hour treatment). Note that NPAA concentrations were selected based on dose-toxicity curves where cell viability was maintained at approximately 80%. Data in **c,d,e,f,g,h,i** are represented as mean ± SEM. n ≥ 3 biological replicates. Student’s *t* test, *p < 0.05; **p < 0.01; ***p < 0.001; n.s., not significant.

**Extended Data Fig. 2.**
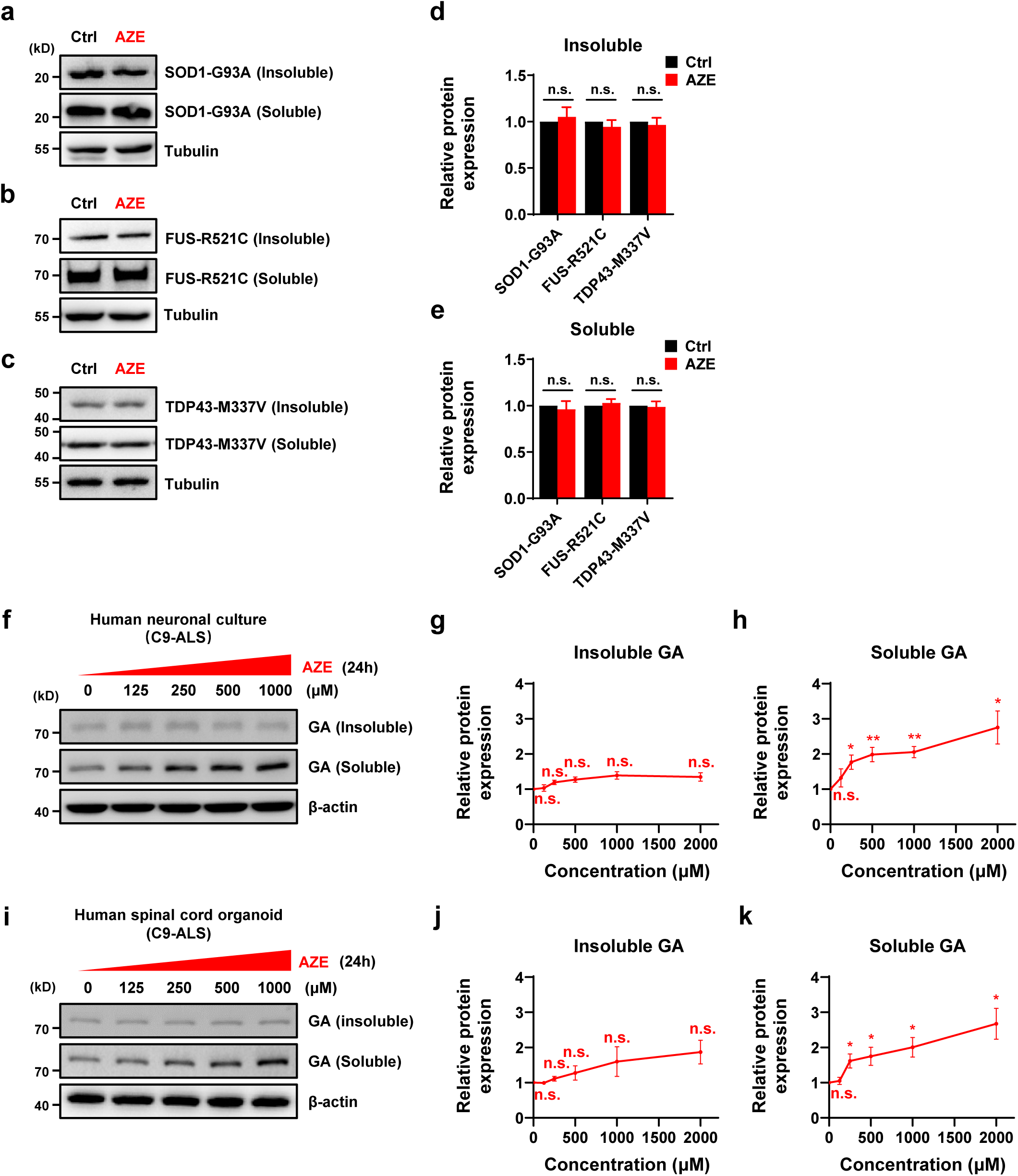
The effect of AZE is specific to poly-GA, not other ALS-associated mutant proteins, related to Fig. 1. **a-e**, Western blot analysis showing the effect of AZE treatment (1000μM, 24 hours) on the expression levels of ALS-associated mutant proteins in HeLa cells. Quantitative analysis was conducted by normalizing both soluble and insoluble protein fractions to Tubulin as a loading control. Note that AZE had no significant effect on SOD1^G93A^, FUS^R521C^, and TDP43^M337V^ levels. **f-k**, Western blot analysis showing the effect of AZE treatment (24 hours) on endogenous poly-GA levels in motor neuron (**f-h**) or spinal cord organoid (**i-k**) cultures derived from C9-ALS patient iPSCs. Quantitative analysis was conducted by normalizing both soluble and insoluble protein fractions to β-actin as a loading control. Note that AZE treatment induced a dose-dependent increase in soluble poly-GA levels, whereas the insoluble forms exhibited minimal changes, suggesting that a critical threshold of poly-GA accumulation may be necessary for the transition into insoluble aggregates. Data in **d,e,g,h,j,k** are represented as mean ± SEM. n ≥ 3 independent experiments. Student’s *t* test, *p < 0.05; **p < 0.01; n.s., not significant.

**Extended Data Fig. 3.**
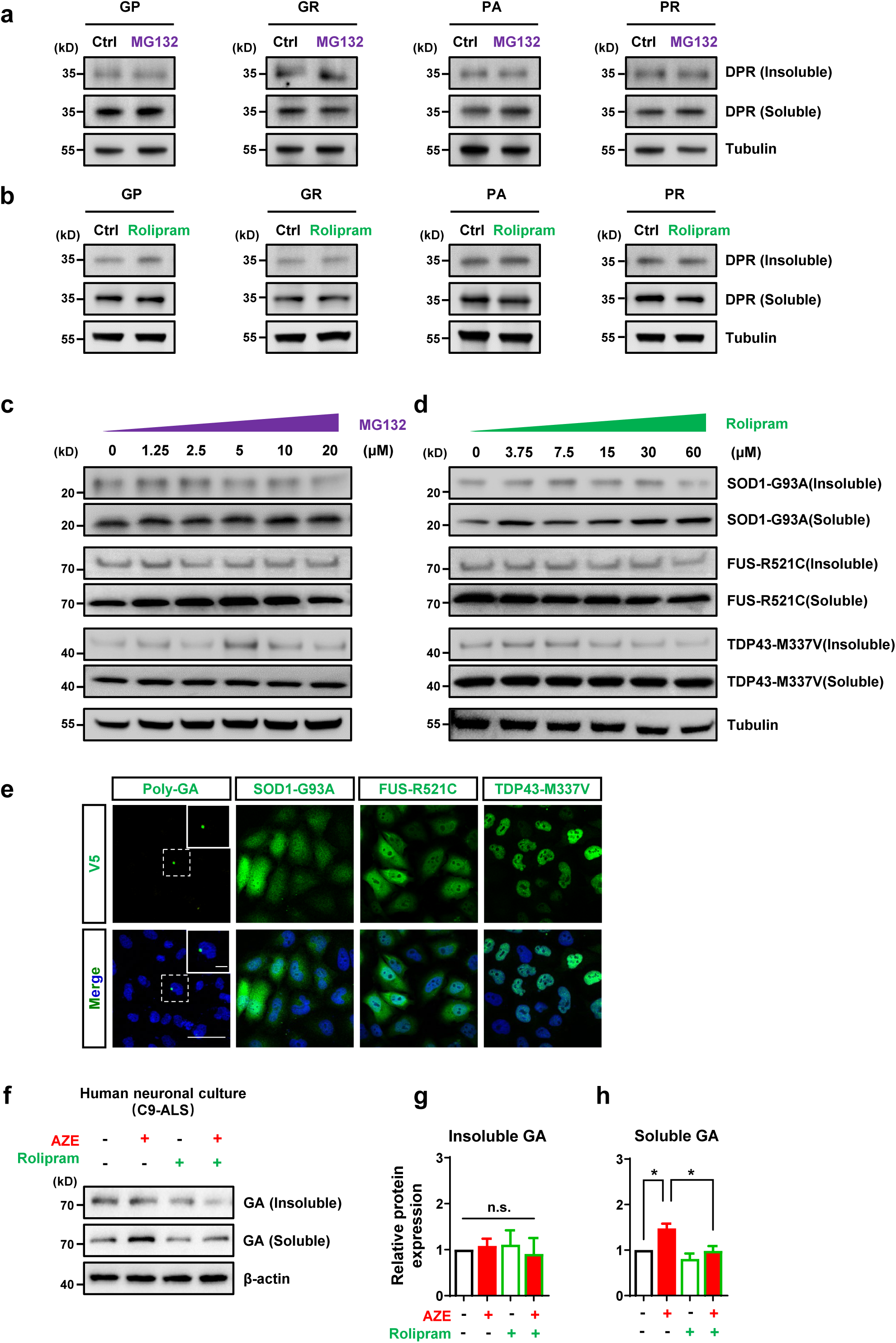
UPS modulation has no effect on levels of other DPRs or ALS-associated mutant proteins, related to Fig. 2. **a**,**b**, Western blot analysis showing the effect of MG132 (10μM, 24 hours) or rolipram (30μM, 24 hours) treatment on DPR levels in HeLa cells transfected with DPR-expressing plasmids: GP(50)-GFP, GR(50)-GFP, PA(50)-GFP, PR(50)-GFP. Note that modulating UPS activity had no detectable effect on these DPR levels. **c**,**d**, Western blot analysis showing the effect of MG132 or rolipram treatment (24 hours) on the expression levels of ALS-associated mutant proteins in HeLa cells. Note that modulating UPS activity had no detectable effect on SOD1^G93A^, FUS^R521C^, and TDP43^M337V^ levels. **e**, Representative images of HeLa cells transfected with plasmids encoding GA(50)-V5, SOD1^G93A^-V5, TDP43^M337V^-V5, and FUS^R521C^-V5. V5 staining is depicted in green, and DAPI staining is shown in blue. Note that poly-GA(50) shows pronounced protein inclusions, a phenotype not shared by other ALS-associated mutant proteins under similar conditions. Scale bar, 50μm. Inset: higher magnification of the white boxed area. Scale bar, 10μm. **f-h**, Western blot analysis showing the AZE-induced upregulation of endogenous poly-GA levels was diminished by rolipram treatment in motor neuron cultures derived from C9-ALS patient iPSCs (AZE, 1000μM; rolipram, 30μM; 24-hour treatment). Quantitative analysis was conducted by normalizing both soluble and insoluble protein fractions to β-actin as a loading control. Note that AZE treatment induced a significant increase in soluble poly-GA levels, whereas the insoluble forms exhibited minimal changes. This observation is consistent with Extended Data Fig. 2f-k. Data in **g,h,** are represented as mean ± SEM. n ≥ 3 independent experiments. Student’s *t* test, *p < 0.05.

**Extended Data Fig. 4.**
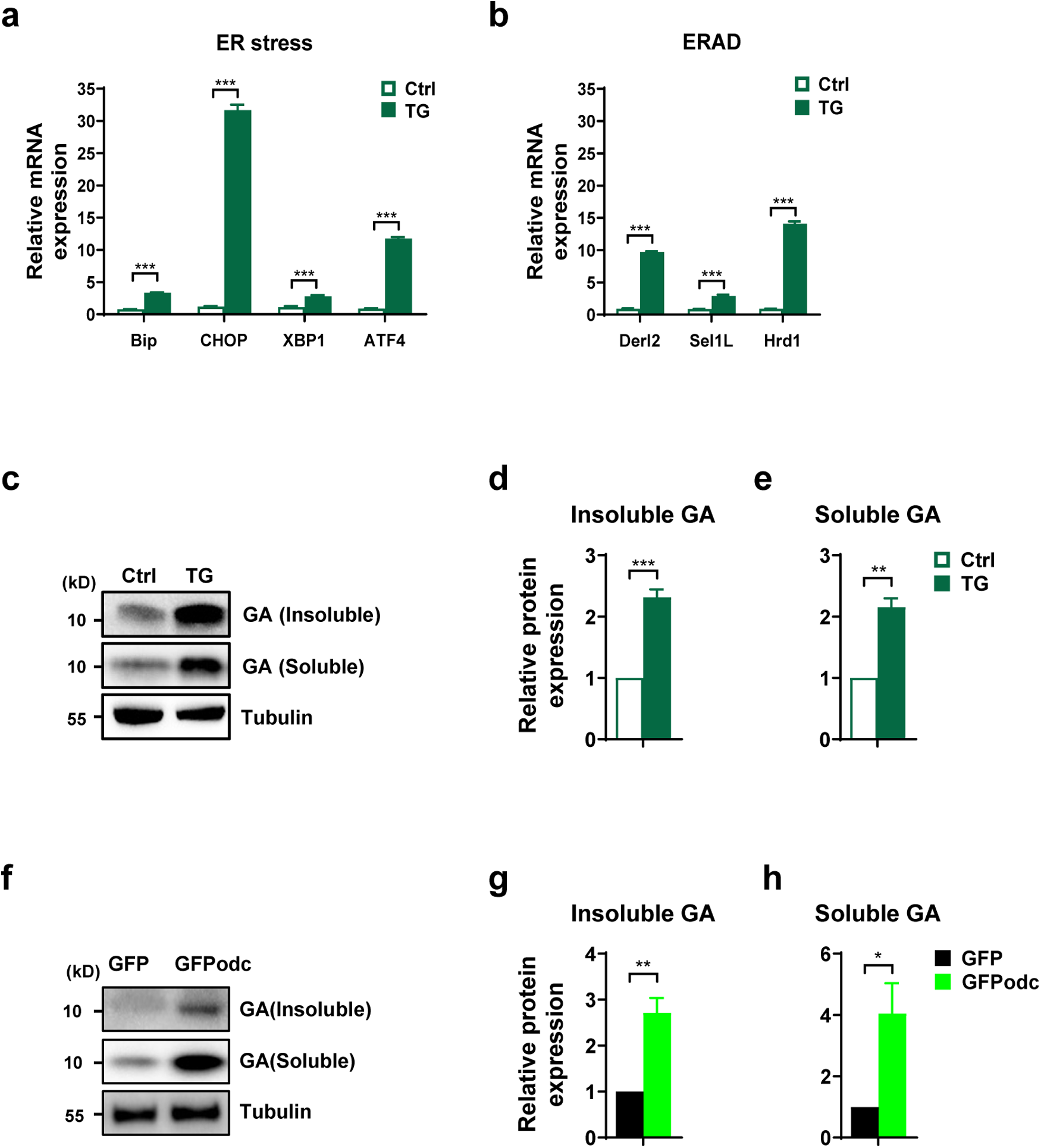
Activation of ERAD or proteasome overload recapitulates AZE-induced poly-GA accumulation, related to Fig. 3. **a**,**b**, qPCR analysis showing that Thapsigargin (TG, 1μM) treatment significantly upregulates the expression of ER stress- and ERAD-related genes in HeLa cells. **c**-**e**, Western blot analysis showing that pharmacological activation of ERAD with thapsigargin (TG, 1μM, 24 hours) is sufficient to induce poly-GA accumulation in HeLa cells transfected with GA(50)-V5-expressing plasmids. Quantitative analysis was conducted by normalizing both soluble and insoluble protein fractions to Tubulin as a loading control. **f**-**h**, Western blot analysis showing that inducing proteasome overload by overexpressing a direct proteasome substrate (GFPodc) induces poly-GA accumulation in HeLa cells expressing GA(50)-V5. Quantitative analysis was conducted by normalizing both soluble and insoluble protein fractions to Tubulin as a loading control. Data in **a,b,d,e,g,h** are represented as mean ± SEM. n ≥ 3 independent experiments. Student’s *t* test, *p < 0.05; **p < 0.01; ***p < 0.001.

**Extended Data Fig. 5.**
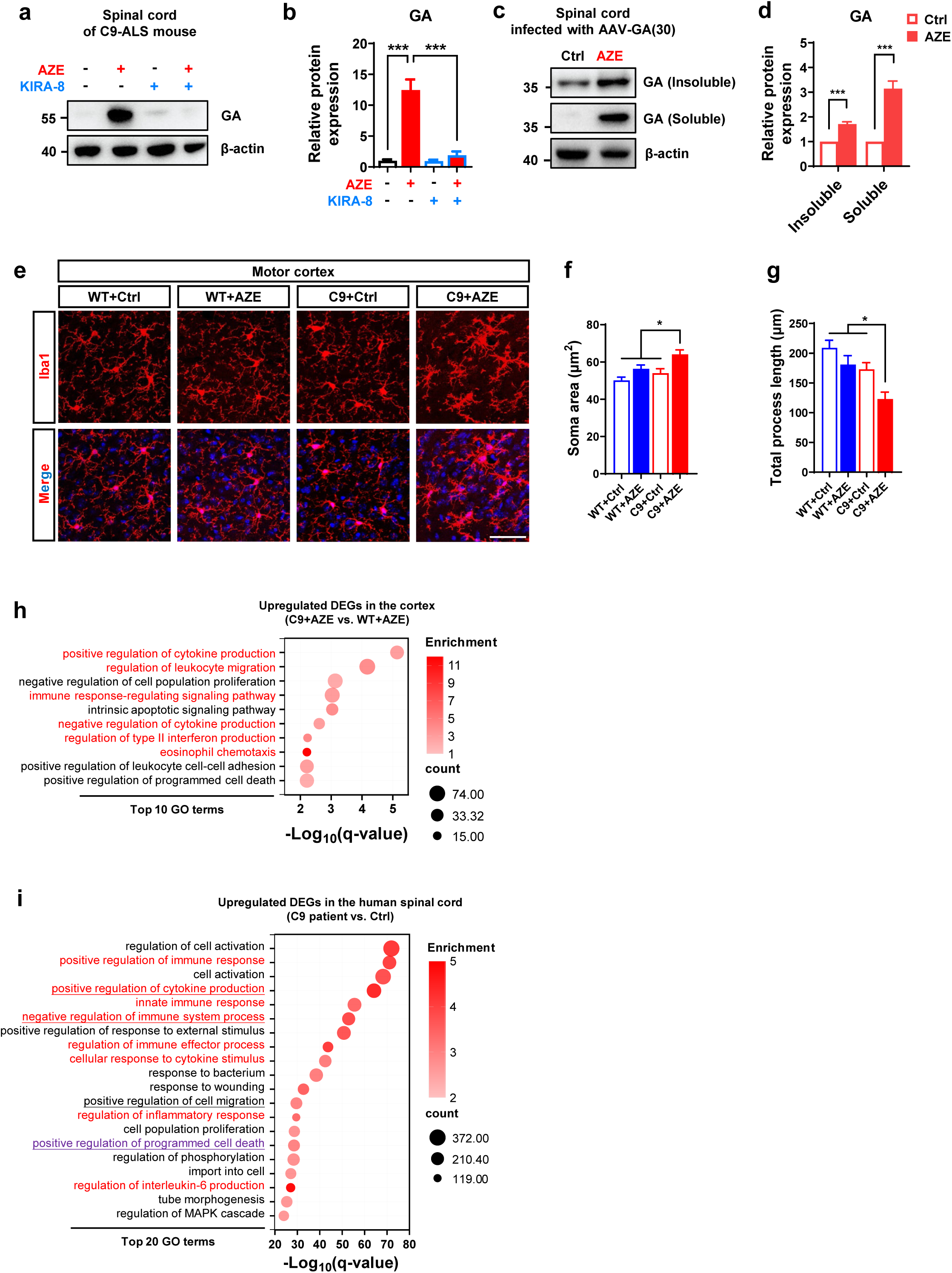
AZE exposure induces neuroinflammation in C9-ALS mice, related to Fig. 4. **a**,**b**, Western blot analysis of spinal cord tissue from C9-ALS mice showing that treatment with the ER stress inhibitor KIRA8 (20 mg/kg, 1 day) attenuates the AZE-induced (350 mg/kg, 1 day) increase in poly-GA levels. Quantitative analysis was conducted by normalizing poly-GA levels to β-actin as a loading control. **c**,**d**, Western blot analysis of spinal cord tissue from mice infected with AAV-GA(30)-GFP. Quantitative analysis was conducted by normalizing poly-GA levels to β-actin as a loading control. **e**, Representative images of microglia (anti-Iba1, red) in the motor cortex from wild-type (WT) and *C9orf72*-humanized mouse models (C9-ALS) treated with AZE (350 mg/kg, i.p., 4 weeks) or vehicle control (saline, i.p., 4 weeks). DAPI staining is shown in blue. Scale bar, 50μm. Note the significant microglial activation in AZE-treated C9-ALS mice, characterized by enlarged cell bodies and decreased process length. **f**,**g**, Quantitative analysis of microglia activation in the motor cortex of different treatment groups, as shown in (**c**). n ≥ 3 mice per group. **h**, Bubble plot of Gene Ontology (GO) analysis for genes upregulated in the cortex of AZE-treated C9-ALS mice versus AZE-treated WT mice (350 mg/kg, i.p., 4 weeks), as determined by RNA sequencing (DEG, differentially expressed gene; fold change ≥ 2; q-value < 0.05). Note that the top 10 enriched GO terms are displayed, with inflammatory response-related terms highlighted in red. **i**, Gene ontology (GO) analysis of genes upregulated in the spinal cords of C9-ALS patients compared to healthy controls, as determined by RNA sequencing (DEG, differentially expressed gene; fold change ≥ 2; q-value < 0.05; data are obtained from Target ALS consortium). Underlined terms represent shared pathways among the top 20 enriched GO terms. Inflammation-related terms are highlighted in red. Cell death-related terms are highlighted in purple. Data in **b,d,f,g** are represented as mean ± SEM. One-way ANOVA, *p < 0.05; ***p < 0.001.

**Extended Data Fig. 6.**
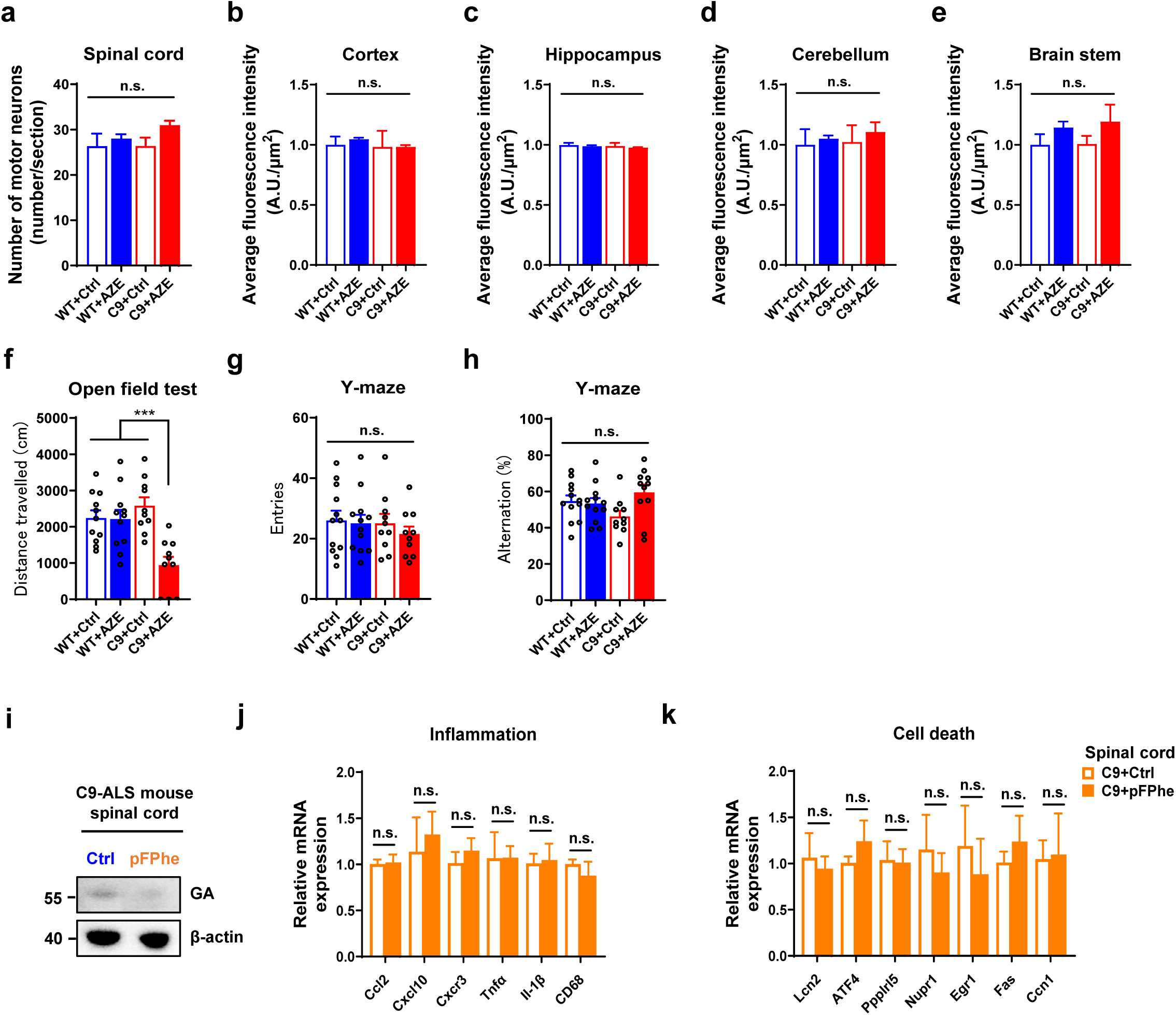
Other toxic NPAA fails to induce comparable pathogenic effects in C9-ALS mice, related to Fig. 5. **a**, Bar graph showing motor neuron counts in the ventral horn of spinal cords from WT and C9-ALS mice treated with AZE (350 mg/kg, i.p., 4 weeks) or vehicle control (saline, i.p., 4 weeks). Motor neurons were labeled using anti-CHAT immunostaining. No significant changes in motor neuron counts following AZE treatment in C9-ALS mice compared to control groups. **b-e**, Bar graph showing no significant changes in the neuronal counts across various brain regions of WT or ALS-C9 mice after AZE treatment (350 mg/kg, i.p., 4 weeks). Neurons were labeled using NeuroTrace staining. A.U., Arbitrary Unit. **f**, Open field test of WT or C9-ALS mice treated with AZE (350 mg/kg, i.p.) or saline for 4 weeks. n ≥ 10 mice per group. **g**,**h**, Y-maze test of wild-type (WT) and *C9orf72*-humanized mouse models (C9-ALS) treated with AZE (350 mg/kg, i.p.) or vehicle control (saline, i.p.) for 4 weeks. n ≥ 10 mice per group. **i**, Western blot analysis of spinal cord tissue from C9-ALS mice treated with pFPhe (350 mg/kg, i.p., 4 weeks) or saline control showing that pFPhe (a toxic NPAA identified in our initial screen) does not increase poly-GA levels. **j**,**k**, qPCR analysis of spinal cord tissue from C9-ALS mice shows that pFPhe treatment (350 mg/kg, i.p., 4 weeks) does not significantly upregulate neuroinflammation- or cell death-related genes (n ≥ 3 mice per group). Data in **b,c,d,e,f,g,h,j,k** are represented as mean ± SEM. One-way ANOVA, ***p < 0.001; n.s., not significant.

**Extended Data Fig. 7.**
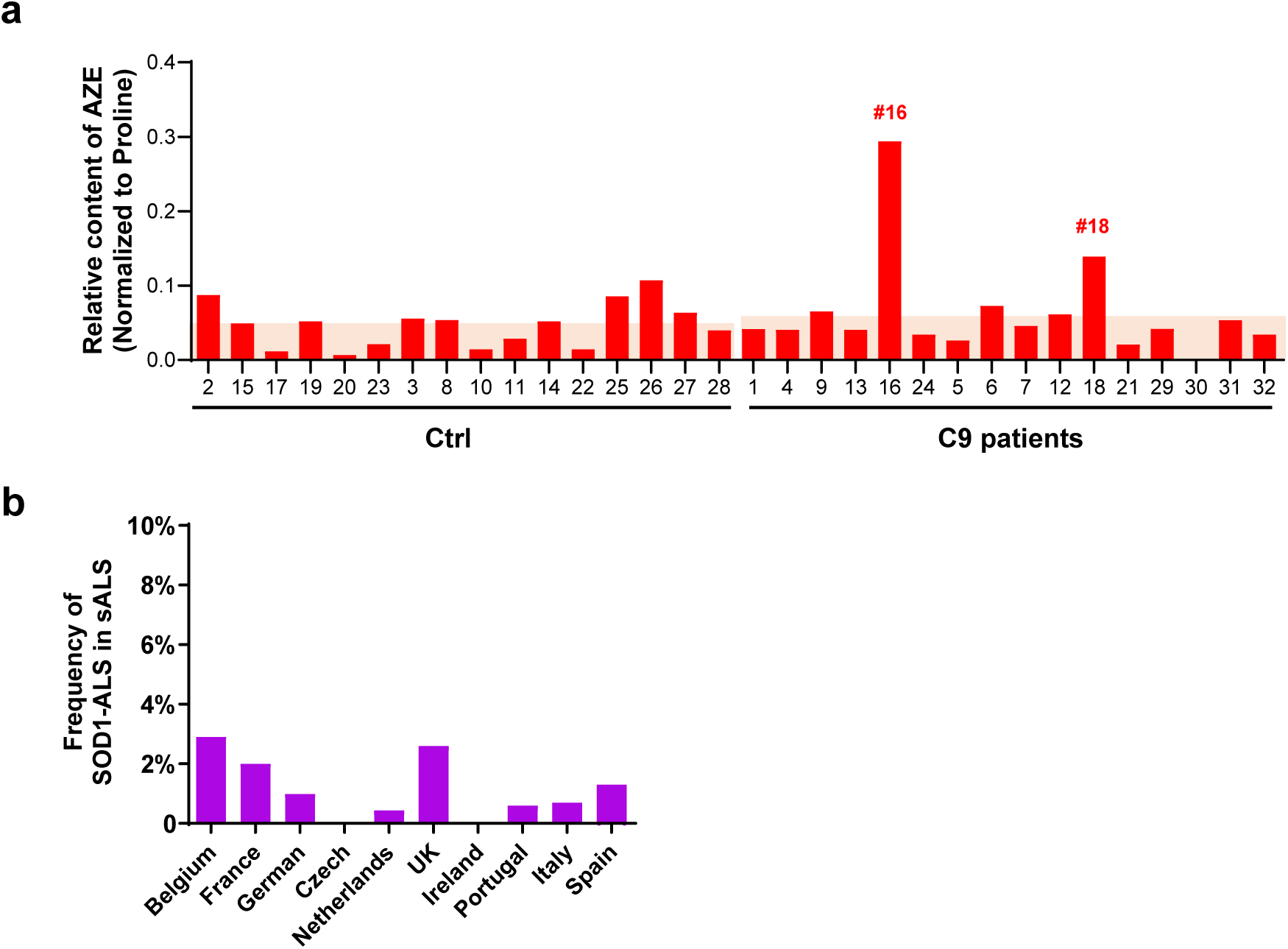
AZE accumulation in postmortem human neural tissues, related to Fig. 7 and 8. **a**, Bar graph showing the relative abundance of AZE (normalized to proline) in spinal cord or cerebellum from C9 patients and normal controls obtained from Netherlands Brain Bank. See Supplementary Table 4 for sample details. The red background indicates the average AZE levels in C9 patients and normal controls, respectively. **b**, The proportion of SOD1-ALS cases among sporadic ALS cases in European countries. Countries were included based on geographic proximity and the availability of reliable national ALS registry data. See Supplementary Table 4 for details.

## Supplemental Table Titles

**Table S1. RNA sequencing analysis of mouse and human neural tissues, related to Fig. 4**

**Table S2. qPCR Primer and NPAA information, related to Fig. 1**, **Fig. 3**, **Fig. 4**, **Fig. 5, and Fig. 7**

**Table S3. Brain Sample Identification and Neuropathological Characteristics, related to Fig. 7**

**Table S4. Sugar beet consumption and the incidence of sporadic C9- and SOD1-ALS cases, related to Fig. 8**

## METHOD DETAILS

### Mice

The following strains of mice were used in this study: wild-type C57BL/6J (Salccas), *C9orf72* transgenic mice (JAX #023099 and #029099; most of experiments were performed using #023099, and parallel experiments using stock #029099 yielded consistent results), *SOD1^G93A^* transgenic mice (JAX #002726).

Both male and female mice were used in this study. All animal studies and experimental procedures were approved by the Animal Care and Use Committee of the animal facility at Zhejiang University. Animals were housed in groups on a 12h light-dark cycle with free access to food and water.

### Human brain tissues

Paraffin-embedded human spinal cord/cerebellum tissues from 16 C9-FTD patients and 16 nondemented controls were obtained from the Netherlands Brain Bank, which supplies postmortem specimens from clinically well documented and neuropathologically confirmed cases (Table S2). All of the material was collected from donors from whom a written informed consent for autopsy, and the use of the material and clinical information for research purposes is available in the Netherlands Brain Bank.

### Construct

The expression plasmids used in this study are as follows: GA(50)-V5, GP(50)-V5, GR(50)-V5, PA(50)-V5, PR(50)-V5, GA(50)-GFP, GP(50)-GFP, GR(50)-GFP, PA(50)-GFP, PR(50)-GFP, SOD1^G93A^-V5, TDP43^M337V^-V5, FUS^R521C^-V5 and GFPodc. Among them, the plasmids GA(50)-V5, GP(50)-V5, GR(50)-V5, PA(50)-V5, and PR(50)-V5, GA(50)-GFP, GP(50)-GFP, GR(50)-GFP, PA(50)-GFP, and PR(50)-GFP were commercially synthesized and cloned into the pN1 vector backbone by Tsingke Biotechnology Co., Ltd. (Beijing, China). For the generation of SOD1^G93A^, TDP43^M337V^, and FUS^R521C^ expression plasmids, the corresponding wild-type sequences were initially PCR-amplified from a mouse whole-genome cDNA library. Site-directed mutagenesis was performed using the Q5 Site-Directed Mutagenesis Kit (New England Biolabs, E0554) to introduce specific mutations. The resulting mutant sequences were subsequently cloned into the pcDNA6 vector backbone using the ClonExpress II One Step Cloning Kit (Vazyme Biotech, C112-01). The plasmid GFPodc was commercially synthesized and cloned into the pQCXIP vector backbone by Tsingke Biotechnology Co., Ltd. (Beijing, China). All constructs were verified through sequencing to ensure sequence accuracy and proper orientation.

### Cell Culture and Transfection

HeLa or NSC34 cells were cultured at 37 °C with 5% CO_2_ in Dulbecco’s modified Eagle’s media (Gibco, 11995) supplemented with 10% fetal bovine serum (Gibco, 10091130) and 1% penicillin/streptomycin (Gibco, 15140-122). Cells were transfected at 80% confluence with Lipofectamine 2000 (Thermo Fisher Scientific, 11668019) according to the manufacturer’s instructions.

### Primary Culture of Spinal Cord Neurons

Primary spinal cord neurons were isolated from the developing spinal cords of E12.5 mouse embryos as previously described.^59^ Briefly, at the desired developmental stage, spinal cords were microdissected with open-book preparation in ice-cold Neurobasal medium (Gibco, 21103-049). Tissue dissociation was performed by incubating the spinal cords in 0.25% trypsin-EDTA (Gibco, 25200-072) at 37 °C for 11-13 minutes. Dissociated neurons were plated onto coverslips pre-coated with poly-D-lysine (PDL; Beyotime, ST508) at 100 μg/mL and laminin (Sigma, L-2020) at 20 μg/mL. Cells were maintained in Neurobasal medium supplemented with 2% B27 supplement (Gibco, 17504044), 2 mM L-glutamine (Invitrogen, 25030-081), 1% penicillin-streptomycin (Gibco, 15140-122), and 50 ng/mL brain-derived neurotrophic factor (BDNF; PeproTech, 450-02). Cultures were incubated at 37 °C in a humidified atmosphere containing 5% CO2. After overnight culture, neurons were treated with various NPAAs at specified concentrations for an additional 24 hours.

### iPSC-derived Motor Neuron Differentiation

iPSCs were obtained from Cedars-Sinai or generated from C9-ALS patients and healthy controls as previously described^29^. Consistent results were observed across multiple cell lines (n ≥ 3 per group). Motor neuron differentiation was performed according to previously described methods with modifications^60^. Briefly, iPSCs were dissociated using Dispase (1 mg/ml; STEMCELL Technologies, 07923) and passaged at a 1:6 ratio onto Matrigel (Corning, 354277)-coated plates. After 24 hours, the PSC medium was replaced with neural induction medium consisting of a 1:1 mixture of DMEM/F12 (Corning, SH30023.FS) and Neurobasal medium (Gibco, 21103049). The medium was further supplemented with 0.5×N2 (Gibco, 17502048), 0.5×B27 (Gibco, 17504044), 0.1 mM ascorbic acid (MCE, HY-B0166), 1×GlutaMAX (Gibco, 35050061), 1×penicillin/streptomycin (Corning, SV30010), 3μM CHIR99021 (MCE, HY-10182), 2μM DMH1 (MCE, HY-12273), and 2μM SB431542 (SELLECK, S1067). Medium changes were performed every other day. After 6 days of induction, iPSCs differentiated into neuroepithelial (NEP) cells.

For motor neuron progenitor (MNP) differentiation, NEP cells were dissociated with Dispase (1 mg/ml) and replated at a 1:6 ratio in neural induction medium supplemented with 0.1μM retinoic acid (RA; MCE, HY-14649), 0.5μM purmorphamine (Pur; MCE, HY-15108), 1μM CHIR99021, 2μM DMH1, and 2μM SB431542. Following 6 days of differentiation, cells expressed the MNP marker OLIG2. MNPs were expanded in medium containing 3μM CHIR99021, 2μM DMH1, 2μM SB431542, 0.1μM RA, 0.5μM Pur, and 0.5μM valproic acid (VPA, MCE; HY-10585), with weekly passaging at 1:6 ratio using Dispase (1 mg/ml). At this stage, cells can be cryopreserved in freezing medium (DMEM/F12, 10% fetal bovine serum, 10% DMSO) and stored in liquid nitrogen for future use.

For motor neuron differentiation, MNPs were dissociated with Dispase (1 mg/ml) and cultured in suspension using neural induction medium supplemented with 0.5μM RA and 0.1μM Pur. Medium changes were performed every other day. After 6 days, cells expressed the motor neuron marker MNX1. Motor neurons were then dissociated into single cells using Accutase (Sigma-Aldrich, A6964) and plated onto Matrigel-coated plates. For terminal maturation into choline acetyltransferase-positive (CHAT^+^) motor neurons, cells were maintained for 10 days in medium containing 0.5μM RA, 0.1μM Pur, and 0.1μM Compound E (MCE, HY-14176). The successful differentiation at each stage was verified by immunostaining with stage-specific markers, as indicated.

### Spinal Cord Organoid Differentiation

Spinal cord organoids were differentiated from iPSCs following established protocols with modifications^61^. Briefly, one day prior to differentiation initiation, approximately 4,000 iPSCs were seeded into each well of a low-adhesion U-bottom 96-well plate (Corning, 3799) in mTeSR1 medium (STEMCELL Technologies, 100-0276) supplemented with 10μM Y-27632 (Selleck, S1049). Embryoid body (EB) formation was initiated by replacing the medium with N2/B27 neural induction medium, consisting of a 1:1 mixture of DMEM/F12 (Corning, SH30023.FS) and Neurobasal medium (Gibco, 21103049) supplemented with 0.5× N2 supplement (Gibco, 17502048), 0.5× B27 supplement (Gibco, 17504044), 1× GlutaMAX (Thermo Fisher Scientific, 35050061), and 1 × penicillin/streptomycin (Thermo Fisher Scientific, 15140122). The medium was further supplemented with 2μM DMH1 (MCE, HY-12273), 10μM SB431542 (SELLECK, S1067), and 3μM CHIR99021 (MCE, HY-10182). Medium changes were performed every other day.

After 4 days of differentiation, EBs were transitioned to caudal neural patterning by replacing the medium with N2/B27 medium supplemented with 0.1μM RA (MCE, HY-14649) and 0.5μM pur (MCE, HY-15108). Medium changes continued every other day. Following 5 days of differentiation under these conditions, EBs contained OLIG2+ motor neuron progenitors.

For terminal differentiation, the medium was replaced with N2/B27 medium supplemented with 0.1μM Compound E (MCE, HY-14176). After 16 days of differentiation, EBs contained CHAT+ motor neurons, representing mature spinal cord organoids. Organoids were maintained with medium changes every other day and could be cultured for extended periods with appropriate maintenance.

### Cell Viability Assay

Cell viability was assessed using the CellTiter-Glo® Luminescent Cell Viability Assay kit (Promega, G7571) according to the manufacturer’s protocol. Briefly, the assay buffer and substrate powder were thawed and equilibrated to room temperature. The substrate was reconstituted in buffer to prepare the enzyme/substrate working solution. Cells (primary cultured spinal cord neurons, iPSC-derived motor neurons, or NSC-34 cells) were cultured in 96-well plates and allowed to equilibrate to room temperature for 30 minutes. An equal volume of working solution was added to each well relative to the culture medium volume (typically 100 μL each). Plates were mixed on an orbital shaker (200 rpm) for 5 minutes to ensure complete cell lysis, followed by a 10-minute incubation at room temperature to stabilize the luminescent signal. The lysate was then transferred to an opaque 96-well plate, and luminescence was measured using a microplate reader at 37 °C.

The hazard ratio was calculated from cell viability assay data using the following approach: First, the percentage of viable cells in each experimental group was normalized to untreated controls. The hazard value was then calculated as (1 - relative viability). Finally, the hazard ratio was determined by dividing the hazard value of C9-ALS samples by the corresponding value from WT controls.

### RNA Isolation and qPCR Analysis

Total RNAs were extracted from cells or tissues using the Easy RNA Kit (Easy-do Hangzhou, DR0401050). Reverse transcription was performed using the HiScript III RT SuperMix for qPCR kit (Vazyme, R323-1) according to the manufacturer’s instructions. Quantitative PCR was conducted using Hieff® qPCR SYBR Green Master Mix (Yeason, 11202ES50) on a QuantStudio real-time PCR system. Gene expression levels were normalized to internal control genes. Primer sequences used in this study are provided in Table S2.

### Viral Delivery

To achieve Poly-GA expression in spinal motor neurons, we delivered rAAV-hsyn-GA(30)-GFP via intramuscular injection into the mouse hindlimb, enabling retrograde transduction of motor neurons. The virus was produced by PackGene Biotech Co., Ltd. (Guangzhou, China) with a titer of 1×10¹³ genome copies per mL. At postnatal day 4 (P4, ±1 day), WT pups were anesthetized on ice and injected with the virus. A total of 2 μL of virus was administered via multiple injections into various hindlimb muscles using a Hamilton syringe. At 8 weeks of age, mice were treated with either AZE (350 mg/kg, 1 day) or vehicle (saline, 1 day). Spinal cord proteins were subsequently extracted for experimental analysis.

### Drug treatment

The NPAA library was purchased from MCE (see Supplemental Table 2 for details). Each compound was dissolved in water or DMSO and applied to NSC-34 cells at a final concentration of 5mM. Cell viability was assessed after 96 hours of treatment.

Neuronal cultures or HeLa cells transfected with specified plasmids were treated with the following compounds for 24 hours: L-Azetidine-2-carboxylic acid (AZE; 125-2000μM; Macklin, L801755), p-fluorophenylalanine (pFPhe; 10-1500μM; Macklin, L809948), m-tyrosine (mTyr; 100-20000μM; Macklin, L863998), L-3,4-dihydroxy-phenylalanine (L-DOPA; 10-160μM; Macklin, L807435), and L-canavanine (CAN; 100-16000μM; Macklin, L918681). MG132(1.25-20μM, MedChemExpress, HY-13259), rolipram (3.75-30μM; MedChemExpress, HY-16900), Thapsigargin (TG, 1μM; MedChemExpress, HY-13433) or KIRA8(0.1μM; MedChemExpress, HY-114368). All compounds were prepared according to manufacturers’ instructions and diluted in appropriate vehicle solutions.

For *in vivo* experiments, C9-ALS mice and WT controls were divided into two groups, respectively. The treatment group received intraperitoneal injections of AZE (350 mg/kg dissolved in sterile saline), pFPhe (350 mg/kg dissolved in sterile saline), TG (1mg/kg dissolved in sterile saline) or KIRA8 (50mg/kg dissolved in vehicle (3% ethanol + 7% Tween-80 + 90% saline)) while the control group received equivalent volumes of vehicle alone. Injections were administered daily for the indicated period.

### Protein Extraction and Western Blot Analysis

Soluble proteins were extracted using RIPA buffer (Fudebio, FD011) supplemented with protease inhibitor cocktail (Bimake, B14002). Tissues or cells were homogenized in lysis buffer and incubated on ice for 30 minutes. Lysates were centrifuged at 12,000 × g for 15 minutes at 4 °C, and the supernatant containing soluble proteins was collected. For insoluble protein extraction, the remaining pellet was resuspended in 8 M urea buffer: 10mM Tris pH 8.0 (Sterile, ST774), 100mM NaH_2_PO_4_ (Hushi Shanghai, 13472-35-0), 8 M urea (Sangon Biotech, A600148-0500) with protease inhibitors. The suspension was vortexed vigorously and centrifuged at 12,000 × g for 15 minutes at 4 °C. Both soluble and insoluble protein fractions were heat-denatured at 95 °C for 5 minutes in loading buffer before analysis.

Proteins were separated by SDS-PAGE and transferred to PVDF membranes. Membranes were blocked with 5% non-fat milk in TBST for 1 hour at room temperature and incubated overnight at 4 °C with the following primary antibodies: anti-GA (Millipore, MABN889); anti-V5 (CST, 13202s); anti-GFP (ProteinFind, HT801-02); anti-ubiquitin (Santa Cruz, sc-166553); anti-SOD1 (proteintech, 10269-1-AP-1); anti-β-actin (Proteintech, 66009-1-Ig); and anti-β-tubulin (Proteintech,10094-1-AP).

### Immunofluorescence staining

For general immunofluorescence staining, cultured cells or tissues were fixed in 4% paraformaldehyde (PFA) at 4 °C. Fixed cells or tissue sections were permeabilized and blocked with PBS containing 1% bovine serum albumin (BSA) and 0.1% Triton X-100 for 1 hour at room temperature. Samples were then incubated overnight at 4 °C with the following primary antibodies: anti-V5 (1:1000; Cell Signaling Technology, 13202S), anti-Chat (1:500; Sigma-Aldrich, AB144P), and anti-Iba1 (1:1000; Abcam, ab178846). Following three washes with PBS, samples were incubated with appropriate Alexa Fluor-conjugated secondary antibodies (488/594; 1:1000) or Neurotrace 640/660 (Thermo Fisher Cat# N21479) for 1 hour at room temperature. Cell nuclei were counterstained with DAPI (1 μg/mL) for 5 minutes.

### Rotarod test

Motor coordination was assessed with a rotarod apparatus (Economex, Columbus Instruments). Mice were acclimated to the apparatus for 3 minutes at a constant speed of 1.0 rpm. For testing, the rotarod was programmed to accelerate from 0 to 40 rpm at a rate of 0.1 rpm per second. Each mouse underwent at least three consecutive trials with 15-20 minute rest intervals between trials. The latency to fall was recorded for each trial, and the average of three trials was calculated as the final motor performance score.

### Grip Strength Assessment

Grips strength was assessed with a grip strength meter (BIO-GS3, BIOSEB, Pinellas Park, USA) as previously described^59^. Mice were gently held by the base of the tail and allowed to grasp the metal grid with their forepaws. The animal was then pulled horizontally away from the device at a consistent speed until it released the grid. The peak force (in grams) was automatically recorded by the device. Each mouse underwent three consecutive trials. The average of three trials was calculated as the final grip strength value.

### Open Field Test

This experiment was conducted in an open field test chamber (manufactured by Anhui Zhenghua). Mice were acclimated to the testing room for 30 minutes prior to the experiment under standardized lighting conditions. Each mouse was gently placed in the corner of the arena (50 cm length × 50 cm width × 30 cm height), facing the wall, and allowed to explore freely for 15 minutes. Behavior was recorded and analyzed using an automated video tracking system. The following parameters were quantified: (1) total distance traveled (cm), (2) time spent in the central zone (16.6 × 16.6 cm), and (3) distance moved in the central zone. Between trials, the arena was thoroughly cleaned with 70% ethanol to remove olfactory cues and any fecal boli.

### Y-maze Test

Spatial working memory and exploratory behavior were evaluated using the Y-maze test, following established protocols^62^. The Y-maze apparatus, comprised three identical arms (50 cm length × 15 cm width × 30 cm height) arranged at 120° angles from each other. During testing, individual mice were gently placed at the distal end of one arm and allowed to freely explore the maze for an 8-minute session. The test was conducted under standardized conditions without prior training or reinforcement (i.e., no food reward or aversive stimuli).

Behavioral parameters were quantified as follows: an alternation was recorded when the mouse sequentially entered all three arms without repetition. The percentage of spontaneous alternation behavior (SAB), representing working memory performance, was calculated using the formula: SAB (%) = [number of alternations/(total arm entries - 2)] × 100. All behavioral sessions were analyzed to determine objective quantification of locomotor activity and arm entries. Mice failing to meet baseline motor performance thresholds were excluded from the analysis.

### AZE Extraction and Quantification

Tissue or food samples were weighed and homogenized in 500 μL distilled water using a mechanical homogenizer (Tiangen, OSE-50). After vortexing for 1 minute, 500 μL methanol was added to the homogenate. The mixture was vortexed thoroughly and centrifuged at 12,000 × g for 10 minutes at 4 °C. The supernatant was collected for LC-MS/MS analysis.

Quantification of AZE was performed using an Agilent 6460 triple quadrupole mass spectrometer (Agilent Technologies, USA) equipped with an electrospray ionization (ESI) source. Chromatographic separation was achieved using a BEH HILIC column (1.7μM, 2.1 × 100 mm) with the following mobile phases: (A) 20mM ammonium acetate (containing 0.1% formic acid) and (B) acetonitrile:methanol (90:10, v/v). Gradient elution was performed at a flow rate of 0.3 mL/min. The mass spectrometer was operated in positive ion multiple-reaction monitoring (MRM) mode with the following parameters: nebulizer pressure at 45 psi, drying gas flow at 5 L/min, sheath gas flow at 11 L/min, and sheath gas temperature at 350 °C. Data acquisition and processing were performed using Agilent Mass Hunter Workstation software.

To ensure measurement consistency, proline was used as an internal control due to its structural similarity and comparable physicochemical properties to AZE. Proline levels were measured concurrently with AZE in all samples, and AZE concentrations were normalized to proline content.

To account for potential alterations in AZE detection resulting from paraffin embedding, we compared AZE levels in mouse tissues before and after paraffin embedding. Fresh mouse tissues from AZE-injected animals were divided into two aliquots: one processed for immediate AZE quantification and the other subjected to paraffin embedding using standard protocols before AZE quantification. The ratio of AZE concentrations in paraffin-embedded versus fresh tissues was calculated and applied as a correction factor to adjust AZE measurements in human paraffin-embedded samples from the Netherlands Brain Bank. All reported brain tissue AZE concentrations in the figures have been normalized using this correction method.

### Sugar Beet Juice Preparation and Administration

Fresh sugar beet roots were thoroughly washed with water and cut into small pieces. The beet pieces were processed using a commercial juicer (SUPOR, TJE06A-400), and the resulting juice was filtered through gauze to remove particulate matter. The filtered juice was aliquoted into sterile tubes and stored at −20 °C until use.

For oral gavage administration, juice aliquots were thawed and equilibrated to room temperature (22-25 °C) to minimize gastrointestinal distress in the mice. A 12-gauge stainless steel gavage needle was used to administer at least 0.4 mL of beet juice per mouse.

### Matrix-Assisted Laser Desorption/Ionization Mass Spectrometry (MALDI-MS) Imaging

MALDI-MS was performed to image metabolite profiles in tissue samples. A matrix solution of 2,5-Dihydroxybenzoic acid (10 mg/mL in 80% ACN (Sigma, 1000292500), 10% MeOH (Sigma, 1060352500), 10% H_2_O and 0.1% TFA (Sigma, 302031-100ML)) was sprayed onto the surface of mouse spinal cord sections using an HTX TM-Sprayer M3 Bundle (HTX Technologies, SP131P924HR). The flow rate of the matrix solution was 100 μL/min. The nozzle temperature and nitrogen gas pressure were set to 70 °C and 10 psi, respectively. The track spacing and velocity of the spray solvent were set to 3 mm and 1,200 mm/min, respectively.

MALDI-MS imaging was performed on a timsTOF fleX MALDI-2 (Bruker Daltonics, Germany) in the Positive ion mode with 400 laser shots per pixel and an interlaser pulse delay of 10 μs controlled by Bruker timsControl software (version 3.0). Transfer settings were 160 V peak-to-peak (Vpp; funnel 1 radio frequency, abbreviated as RF), 160 Vpp (funnel 2 RF), and 160 Vpp (multipole RF). Focus pre-time-of-flight transfer time was set to 45 μs and the pre-pulse storage to 5 μs. The quadrupole ion energy was 10.0 eV with a low mass of m/z = 50. The collision cell energy was 10.0 eV with collision RF at 180 Vpp. All of the spectra were acquired in the m/z range of 20–350 using a 150 × 150μM^2^ pixel size for tissue analysis. Calibration of the instrument was carried out before every measurement with Tuning Mix.

### RNA-seq Data Processing and Differential Expression Analysis

RNA-seq data were obtained from the spinal cord tissues of wild-type C57 mice (control group) and C9orf72 mice (experimental group), both treated with AZE (350 mg/kg, i.p, 4 weeks). Raw sequencing reads were subjected to quality control using FastQC (version 0.11.9). Adapter sequences and low-quality bases were trimmed using Trimmomatic (version 0.39) with the following parameters: SLIDINGWINDOW:4:20 MINLEN:50. The quality-filtered reads were aligned to the mouse reference genome (GRCm39) using STAR aligner (version 2.7.10a). Gene-level read counts were quantified using featureCounts (version 2.0.1). Differential expression analysis was conducted using DESeq2 (version 1.30.1) with the following criteria: absolute log2 fold change > 1 and adjusted p-value (FDR) < 0.05. Volcano plots were generated to visualize the differentially expressed genes (DEGs) between the control and experimental groups.

### Gene Ontology (GO) Enrichment Analysis

To explore the biological functions and pathways associated with the DEGs identified in the mouse spinal cord and motor cortex, Gene Ontology (GO) enrichment analysis was performed using Metascape (http://metascape.org). The DEG lists, including gene symbols, were uploaded to the Metascape platform, which were annotated through the platform with analysis restricted to the Biological Process (BP) domain. The Benjamini-Hochberg procedure was applied to adjust for multiple testing. Enrichment analysis was conducted with the following parameters: a minimum overlap of 5 genes, a q-value cutoff of 0.05, and a minimum enrichment factor of 2. Bubble plots were generated using GraphPad Prism (version 10.4.1) to visualize the enriched GO terms based on their q-values, enrichment factors, and counts (the number of significantly changed genes within each GO term).

To assess the similarity between AZE-treated C9-ALS mice and human ALS patients, we conducted a comparative analysis of their GO enrichment profiles. RNA-seq data from human ALS patients were acquired from the TargetALS consortium and processed using the same analytical pipeline as described for mouse samples^41^. Significant differentially expressed genes (DEGs) in human spinal cord tissues (C9-ALS patients vs. controls) were identified using an adjusted p-value threshold of < 0.05. GO enrichment analysis was performed on both mouse and human DEGs using Metascape with identical parameters as described above. Conserved biological processes were identified through comparative analysis of the mouse and human GO enrichment results, focusing on overlapping terms with consistent directional changes in gene expression.

### Quantification and Statistical Analysis

All quantitative data are presented as mean ± SEM. Statistical analyses were performed using GraphPad Prism 9 software. Differences between two groups were analyzed using either a two-tailed Student’s *t*-test (for normally distributed data) or the Mann-Whitney U test (for non-normally distributed data). Differences between multiple groups were analyzed using Ordinary One-way ANOVA. P-values were corrected for multiple comparisons using Dunnett’s multiple comparisons test for One-way ANOVA. Statistical significance was defined as *p < 0.05, **p < 0.01, and ***p < 0.001. Unless otherwise specified, each experiment was independently replicated at least three times. For animal studies, behavioral experiments were conducted by investigators blinded to genotype and treatment conditions. Mice were randomly assigned to experimental groups, with approximately equal numbers of male and female animals included in each group. No sex-specific differences in disease progression or treatment response were identified. Sample sizes were determined based on previous studies.

In our statistical analysis, the software automatically performs normality tests (e.g., Shapiro-Wilk test) and tests for homogeneity of variances (e.g., Brown-Forsythe test). Based on the outcome of these tests, the appropriate statistical method and corresponding p-value are selected. For comparisons involving multiple groups, if the data passes both normality and equal variance assumptions, we use Ordinary one-way ANOVA followed by a post-hoc test for multiple comparisons. If these assumptions are violated, the non-parametric Kruskal-Wallis test, followed by an appropriate post-hoc test for multiple comparisons, is applied.

